# ZO-1 shuttles between tight junctions and podosomes by riding ERK activation waves during collective cell migration

**DOI:** 10.1101/2024.08.09.607278

**Authors:** Sayuki Hirano, Yohei Kondo, Noriyuki Kinoshita, Tetsuhisa Otani, Mikio Furuse, Naoto Ueno, Kazuhiro Aoki

## Abstract

Collective cell migration is essential in various physiological processes, including embryonic development, wound healing, and cancer metastasis. However, the mechanisms by which individual cells achieve coordinated movement remain elusive. Here, we demonstrate that zonula occludens-1 (ZO-1), a scaffolding protein of tight junctions (TJs), dynamically translocates to form cell-to-extracellular matrix (ECM) adhesion complexes, podosomes, at the basal cell surface during migration. The translocation of ZO-1 is regulated by extracellular signal-regulated kinase (ERK) activity and its interaction with F-actin. ZO-1 at podosomes contributes to traction force generation and cell invasion, while ZO-1 at TJs mediates intercellular propagation of ERK activation, facilitating collective cell migration. These findings elucidate the versatile roles of ZO-1 in cell adhesion and movement, providing insights into the mechanisms coordinating the discrete movements of individual cells into a cohesive and directed collective cell migration.

## Introduction

Collective cell migration constitutes a fundamental mechanism underlying various physiological phenomena, including embryonic development and wound healing^1^. Cells migrate as a cohesive group in response to extracellular environmental stimuli, such as chemical gradients and substrate stiffness^2,3^. Interestingly, it has been reported that certain cells exhibit directional migration only when they are in a collective state, and randomly migrate when in an individual state^4^, suggesting that single-cell migration mechanisms are not sufficient to fully explain collective migration. Collective cell migration also plays a significant role in cancer metastasis, with many types of cancer cells known to invade in a cohesive manner^5^. Therefore, it is critical to elucidate the mechanisms of collective cell migration not only to improve our understanding of developmental processes but also to identify potential therapeutic targets in cancer treatment.

How individual cells that only sense their surroundings can achieve coordinated collective migration on spatial scales larger than themselves remains elusive. Successful collective migration requires mechanisms that enable intercellular communication and that integrate these signals into the regulation of cell movement. Among the structures involved in intercellular communication are adherens junctions, which play a role in the transmission of mechanical tension between cells. It has been reported that downregulation of adherens junction proteins or inhibition of their mechanoresponsive function suppresses directional collective cell migration^3,4,6^. Additionally, extracellular signal-regulated kinase (ERK) has been increasingly recognized as a significant signaling molecule that regulates collective cell migration. We have recently shown that ERK activation is propagated as waves across large numbers of cells during collective migration of cultured cells^7,8^ and mouse ear skin^9^. The intercellular propagation of ERK activation depends on the epidermal growth factor receptor (EGFR) signaling pathway, which is activated by extracellularly released ligands^7,10^ and/or mechanical stimuli from the neighboring cells^11,12^. Although ERK propagation drives collective cell migration by regulating actomyosin contractility, it is still unclear what specific substrates ERK phosphorylates to control cell movement.

Tight junctions (TJs) are one of the two major forms of intercellular adhesion complexes, with adherens junctions being the other. In epithelial cell sheets, TJs are located apically to adherens junctions and serve to tightly connect adjacent cell membranes, restricting the leakage of molecules through the paracellular space^13^. While the role and molecular assembly of TJs in barrier formation is well understood, their function in collective cell migration is largely unknown. Several studies have reported that zonula occludens-1 (ZO-1), a major scaffolding protein of TJs, influences collective cell migration by controlling actomyosin contractility^14–16^. ZO-1 negatively regulates contractile forces at the cell periphery, and its depletion leads to hyper-development of actomyosin cables encircling individual cells, thereby reducing tissue fluidity and inhibiting collective cell migration^14–16^.

Additionally, we have recently suggested a relationship between ZO-1 function and ERK activity; in *Xenopus* embryos, centrifugal forces enhance both ERK activation and ZO-1 phosphorylation^17,18^. These findings led us to hypothesize that ZO-1 controls collective cell migration downstream of ERK through its interaction with actomyosin regulatory networks.

In this study, we report the dynamic behavior of ZO-1 during collective cell migration using a commonly used epithelial cell model, the Madin-Darby canine kidney (MDCK II) cell. In a migrating cell population, ZO-1 transiently moved from TJs to distinct cell–extracellular matrix (ECM) adhesion complexes known as podosomes. This translocation of ZO-1 was regulated by ERK activity and direct interaction with F-actin, contributing to traction force generation and cell invasion. Additionally, ZO-1 that remained at TJs facilitated collective cell migration by mediating intercellular propagation of ERK activity. These findings elucidate the versatile roles of ZO-1 in the regulation of collective cell migration and invasion.

## Results

### ZO-1 translocates from TJs to podosomes during collective cell migration

To explore the role of ZO-1 in collective cell migration, we first examined the behavior of ZO-1 in migrating cell populations using MDCK II (Madin-Darby Canine Kidney II) cells. We established a stable cell line expressing EGFP-tagged ZO-1 on a ZO-1/2 double knock-out (ZO-1/2 dKO) MDCK II background^19^. Unless otherwise specified, MDCK II ZO-1/2 dKO cells were used in subsequent experiments. The cells were first seeded to confined areas of a culture insert and cultured for one day until they formed a confluent monolayer. After that, the insert was removed so that the cells could migrate into free spaces. Prior to the onset of migration, ZO-1 primarily localized at the apical part of the cells and formed TJs (Fig. 1A, B, top panels). To our surprise, after the cells started migration, ZO- 1 dissociated from TJs and relocated to the basal part of the cells, accumulating at the front region of individual cells (Fig. 1A, B, lower panels; Video S1). ZO-1 in the basal-front region of individual cells often assembled into a circular pattern (Fig. 1A, arrowheads) or into an irregular cluster (Fig. 1A, arrows). Furthermore, this translocation of ZO-1 was propagated through the migrating cell population, initially appearing in the leading-edge cells and then progressively in the following cells.

**Figure 1.**
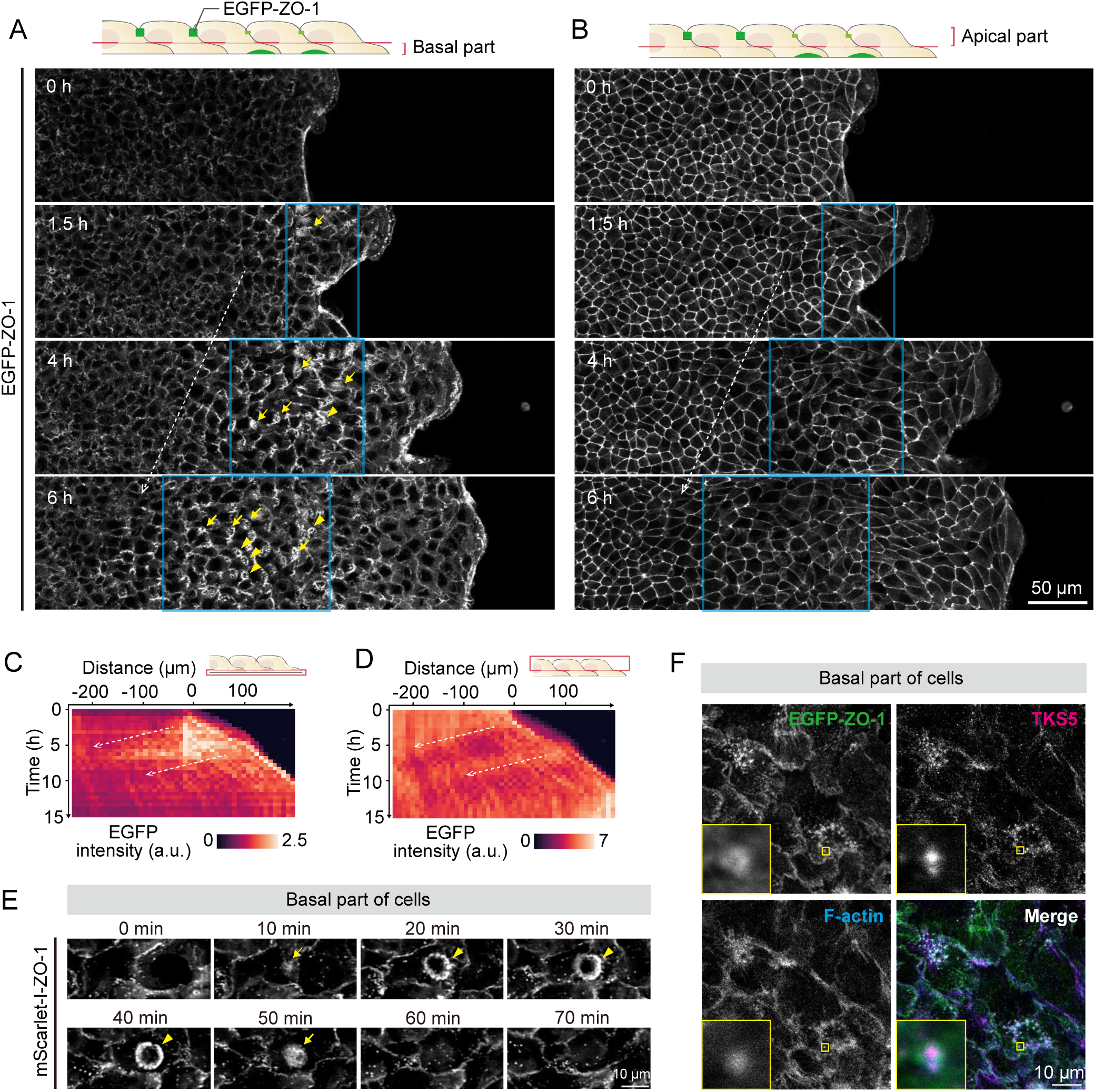
Intercellularly propagating waves of ZO-1 translocation from TJs to podosomes during collective cell migration. (A and B) Time-lapse images of a migrating MDCK II cell population stably expressing EGFP-ZO-1 in the backbone of ZO-1/2 double KO. The basal part (A) and the apical-lateral part (B) of the same cell population are represented. Arrowheads and arrows indicate circular assemblies and irregular clusters of ZO-1, respectively. (C and D) Kymographs of the average fluorescence intensity of panel A and B. (E) Time-lapse images of the basal part of a migrating MDCK II cell population that stably expresses mScarlet-I-ZO-1 in the backbone of ZO-1/2 dKO cells. Arrowheads and arrows indicate circular assemblies and irregular clusters of ZO-1, respectively. (F) Immunostaining images of MDCK II ZO-1/2 dKO/EGFP-ZO-1 cells. Inserts are the magnifications of the yellow boxes. The direction of the migration is from left to right in all images.

Immunostaining confirmed that endogenous ZO-1 also accumulated at the basal-front region of migrating cells (Fig. S1), indicating that this translocation was not due to an artificial effect of EGFP tagging or genome integration. To quantify the localization changes of ZO-1, we measured the fluorescence intensity of EGFP-ZO-1 at the basal and apical part of the cells. The results showed waves of increasing fluorescence intensity at the basal part and also the accompanying waves of decreasing fluorescence intensity at the apical part (Fig. 1C, D). We confirmed that the change in ZO-1 intensity was not simply due to positional changes of membrane components as the cells elongated and flattened during migration by obtaining the ratio of EGFP-ZO-1 to mScarlet-I-KRasCT (KRasCT: the C-terminus of KRas, a plasma membrane localizer) (Fig. S2). These results suggest that ZO-1 plays a distinct role in migrating cell populations at the basal part, apart from its role in TJ formation.

To identify the nature of the basal ZO-1 structure, we closely observed its formation and dissolution process in migrating cell populations (Fig. 1E; Video S2). ZO-1 initially accumulated at the front edge of a cell, followed by the appearance of an irregular cluster at the basal plane of the cell (Fig. 1E, arrow). The ZO-1 cluster then developed to a circular assembly (Fig. 1E, arrowhead), and the cell body moved forward, while the position of the circular assembly remained largely unchanged. Within several tens of minutes to an hour, the circular assembly of ZO-1 dissolved, and ZO-1 again formed an irregular cluster before disappearing from the basal cell surface. The clusters and assemblies of ZO-1 were akin to those of podosomes in their appearance. Podosomes are a type of cell–ECM adhesion complex that promote proteolytic invasion of cells^20^. Each podosome consists of a core containing F-actin, TKS5, and MT1-MMP, and an adhesive ring containing integrins and vinculin. The core contributes to ECM degradation^21,22^ and the generation of protrusive forces against the substrate^23,24^.

The ring allows adhesion to the ECM and generates pulling forces, which enable the oscillatory protrusion of podosomes in coordination with the protrusive forces of the core^25,26^. Podosomes often organize in large numbers, resulting in the formation of higher-order structures such as circular assemblies, belt-like assemblies, and irregular clusters^27–29^. To examine whether the basal ZO-1 structures observed in migrating cells were assemblies of podosomes, we performed immunostaining for the podosome markers TKS5 and F-actin, together with ZO-1. The results showed that ZO-1 was co-localized with the podosome markers in the basal-front region of migrating cells, indicating that the basal ZO-1 structures were assemblies of podosomes (Fig. 1F). Closer observation of individual podosomes revealed that ZO-1 was localized at the periphery of the core consisting of TKS5 and F-actin (Fig. 1F, magnified), suggesting its interaction with adhesion molecules in the adhesive ring of podosomes. Taken together, these results indicate that ZO-1 dissociates from TJs and relocates to podosomes during collective cell migration.

### ZO-1 is recruited to podosomes through the actin-binding region

To elucidate the mechanism by which ZO-1 is recruited to podosomes, we focused on F-actin, one of the major components of podosomes. ZO-1 interacts with F-actin via its actin-binding region (ABR) at the C-terminus^30,31^ (Fig. 2A). This interaction is required for the dynamic remodeling and barrier function of TJs^32,33^. We utilized a mutant ZO-1 that lacks ABR (ZO-1ΔABR^34^) to examine the role of the ABR in the subcellular localization of ZO-1. In a confluent monolayer, The ZO-1ΔABR was mostly localized at TJ regions, like full-length ZO-1, but also partially localized in the cytoplasm (Fig. 2B and 2C). Next, we used 12-O-tetradecanoylphorbol 13-acetate (TPA; also known as phorbol 12- myristate 13-acetate: PMA), which is widely used to induce podosome formation in various cell types through protein kinase C (PKC) activation^35–37^. Several tens of minutes after the addition of TPA, full-length ZO-1 had accumulated in podosomes and cell peripheries on the basal side (Fig. 2D; Video S3). TPA-induced podosomes formed irregular clusters rather than forming circular assemblies as observed in migrating cells. In contrast to full-length ZO-1, ZO-1ΔABR did not efficiently accumulate in podosomes even after TPA addition (Fig. 2D, lower panels; Video S3). The percentage of cells with ZO-1 accumulation in podosomes was significantly lower in ZO-1ΔABR than in full-length ZO-1 (Fig. 2F). Closer observation revealed that ZO-1 was localized at the adhesive ring surrounding the actin core of podosomes (Fig. 2G), as observed in migrating cells. When ZO-1 lacked the ABR, it no longer accumulated in the adhesive ring, although the actin core was formed (Fig. 2H). We also found that the absence of ZO-1 in the adhesive ring of podosomes reduced the recruitment of another ring-forming protein, vinculin (Fig. S3). In the cells expressing full-length ZO-1, vinculin was localized to both of podosomes and focal adhesions, whereas in the cells expressing ZO-1ΔABR, vinculin was localized only to focal adhesions (Fig. S3). These results indicate that ZO-1 is localized to podosomes through its actin-binding region and is involved in the formation of the adhesive ring rather than the actin core of podosomes.

**Figure 2.**
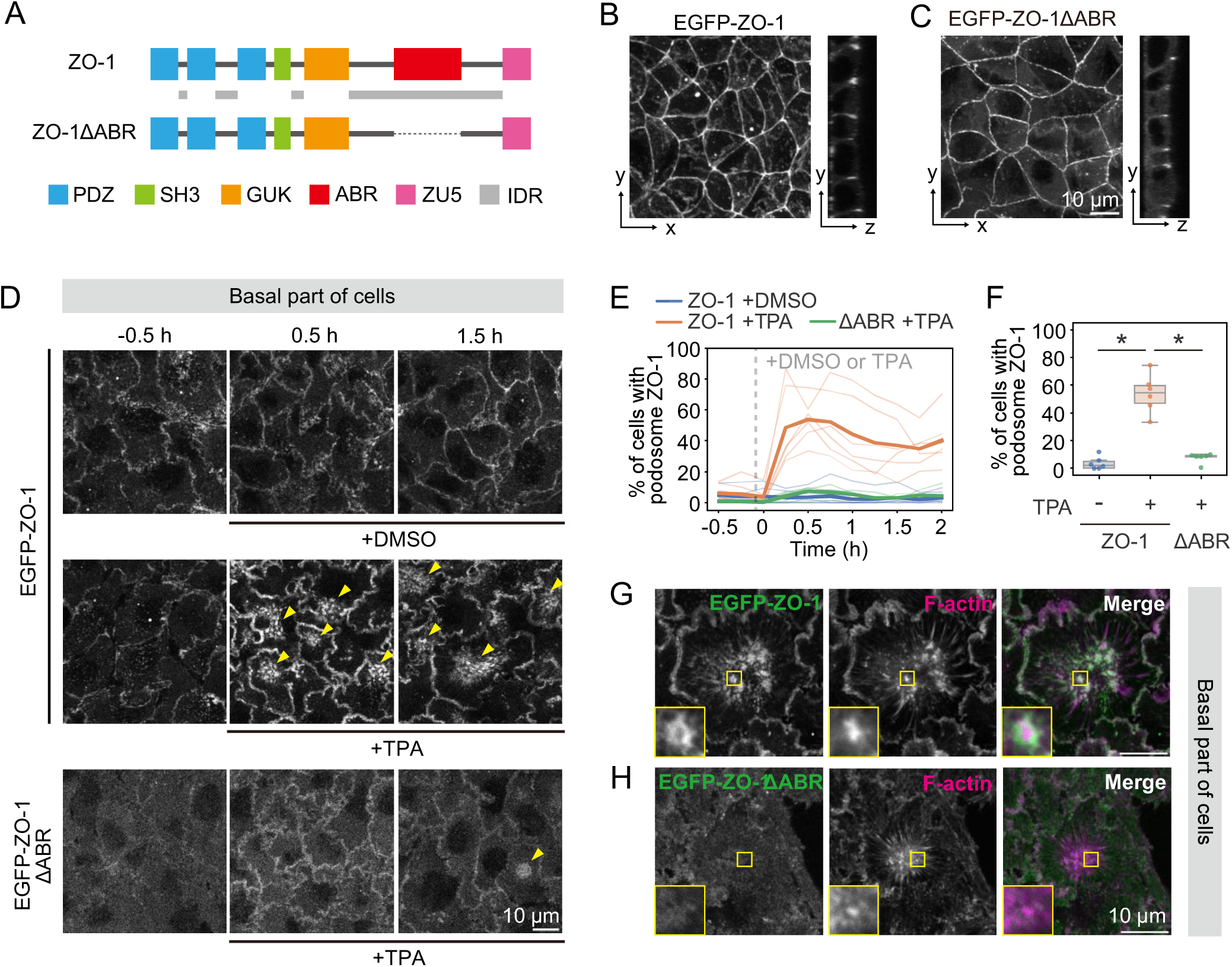
Involvement of the actin-binding region in ZO-1 localization to podosomes (A) Schematic images of ZO-1 and ZO-1ΔABR (actin-binding region). (B and C) Apical (left) and lateral view (right) of MDCK II ZO-1/2 dKO/EGFP-ZO-1 (B) and MDCK II ZO-1/2 dKO/EGFP-ZO-1ΔABR (C). (D) Time-lapse images of the basal part of the cells treated with DMSO or 10 nM TPA in the indicated cells. Arrowheads indicate clusters of podosomes. (E and F) Quantified data of panel D. Data are from three independent experiments. Thin lines and bold lines in panel E indicate the results of each sample (n = 6 for each condition) and their average, respectively. Panel F shows the data 30 min after the drug treatment. Statistical significance was tested by the Kruskal-Wallis test and the Steel–Dwass test. (G and H) Immunostaining images of MDCK II ZO-1/2 dKO/EGFP-ZO-1 (G) and MDCK II ZO-1/2 dKO/EGFP-ZO-1ΔABR (H). The cells were treated with 10 nM TPA 1 h before fixation. Inserts are the magnifications of the yellow boxes.

### ZO-1 translocation to podosomes follows ERK activation during collective cell migration

We next examined the involvement of ERK activity in the translocation of ZO-1 to podosomes during collective cell migration, for the following three reasons: (1) ERK is activated downstream of the PKC pathway and is involved in the formation and the function of podosomes^38,39^. (2) Intercellular propagation of ERK activation is one of the key regulators of collective migration of epithelial cells^7,11^. (3) ERK activation and ZO-1 phosphorylation are simultaneously induced by mechanical stimuli^17^. ERK activity was visualized using a fluorescence resonance energy transfer (FRET)-based ERK activity sensor (EKAREV-NLS^40^; Fig. 3A, upper left). This sensor undergoes a conformational change upon phosphorylation by activated ERK and exhibits an increase in the FRET/CFP ratio corresponding to FRET efficiency (see Materials and Methods for more details). Along with probing ERK activity, we investigated the formation of circular assemblies of podosomes (Fig. 3A, lower left) using U-Net, a deep learning model for semantic segmentation of biomedical images. In this analysis, we considered circular assemblies of ZO-1 as circular assemblies of podosomes, because ZO-1 was colocalized well with the podosome marker TKS5 on the basal cell surface during collective cell migration (Fig. 1F). The model was trained with manually annotated images of circular assemblies of podosomes and used for segmentation of those structures in fluorescence images of mScarlet-I-ZO-1 (Fig. 3A, upper right) (see Materials and Methods for more details). Previous studies have shown that ERK activation is propagated as waves within a migrating epithelial cell population and correlated well with cell deformation^7,11^. In agreement with the observations in these studies, we observed several ERK activation waves in migrating cell populations (Fig. 3B). Initially, ERK activation propagated as a slow wave (Fig. 3B, dashed arrow; 1.1 ± 0.4 μm/min) from the leading edge to the following cell population, and then propagated as several fast waves (Fig. 3B, dotted arrows; 2.1 ± 0.8 μm/min). The formation of circular assemblies of podosomes appeared as propagating waves in the accompanying ERK activation waves (Fig. 3C). A closer look revealed that ERK was activated prior to podosome formation (Fig. 3D, 80 min; Video S4). Several tens of minutes after ERK activation, circular assemblies of podosomes appeared in the front side of well-elongated migrating cells (Fig. 3D, 130 min). During periods of sustained high ERK activity, cells often exhibited oscillatory formation of circular assemblies of podosomes (Fig. 3D, arrowheads). Analysis of the temporal changes in ERK activity and podosome formation revealed multiple peaks in the formation of circular assemblies of podosomes around a peak of ERK activity (Fig. 3E). In addition, when ERK activity decreased, podosome formation was distinctly suppressed. These results suggest that ZO-1 is translocated from TJs to podosomes by ERK activation during collective cell migration.

**Figure 3.**
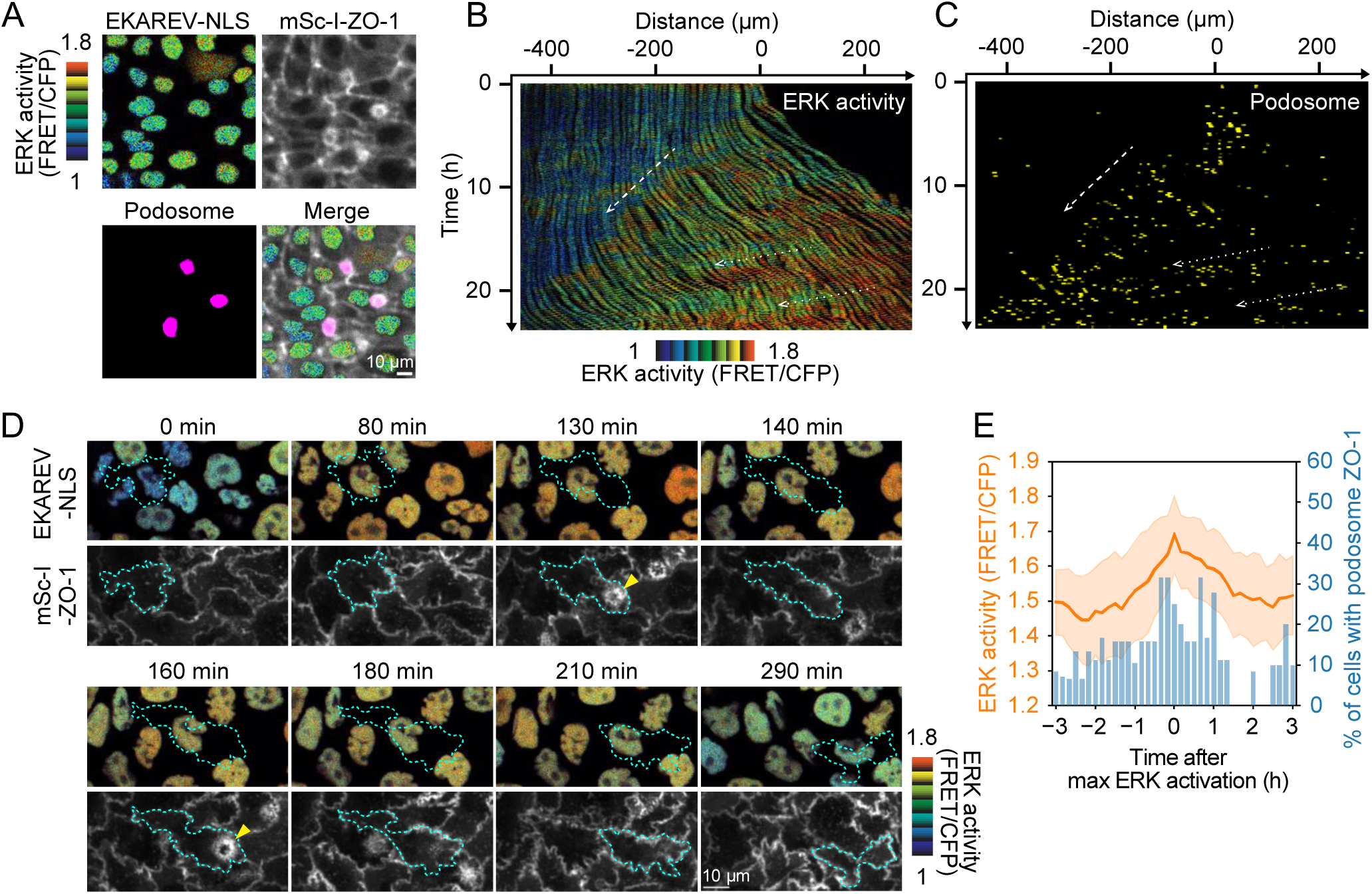
ZO-1 translocation to podosomes following ERK activation waves during collective cell migration (A) Representative images of EKAREV-NLS (FRET/CFP), mScarlet-I-ZO-1 and detected podosome assemblies. A high FRET/CFP ratio indicates high ERK activity. (B, C) Kymographs of ERK activity (B) and podosome assemblies (C) in collectively migrating MDCK II ZO-1/2 dKO/EKAREV-NLS/mScarlet-I-ZO-1 cells. A dashed arrow and dotted arrows indicate a slow wave and fast waves, respectively. (D) Time series of ERK activity and ZO-1 assembly into podosomes during collective cell migration. Arrowheads indicate circular assemblies of podosome ZO-1. Time-course of ERK activity and ZO-1 assembly into podosomes. The time point at which ERK activity (FRET/CFP) in each cell reached its peak was set as time zero. The line and the band of ERK activity indicate average and s.d., respectively. The bars indicate the percentage of cells with circular assembly of podosome ZO-1. N = 20.

### ERK activity controls ZO-1 translocation to podosomes

To determine whether ERK activity affects the translocation of ZO-1 to podosomes, we treated the cells with a MEK inhibitor (MEKi, PD0325901) after inducing podosome formation by TPA. Upon addition of the MEKi, the TPA-induced podosomes immediately disappeared and remained at a low level for at least 1 h, while DMSO treatment caused only a slight reduction in podosome formation (Fig. 4A and 4B; Video S5). The percentage of cells with ZO-1 accumulation in podosomes was significantly lower in the MEKi-treated group compared to the DMSO-treated group (Fig. 4B). This result indicates that ERK activity is necessary for ZO-1 accumulation in podosomes.

**Figure 4.**
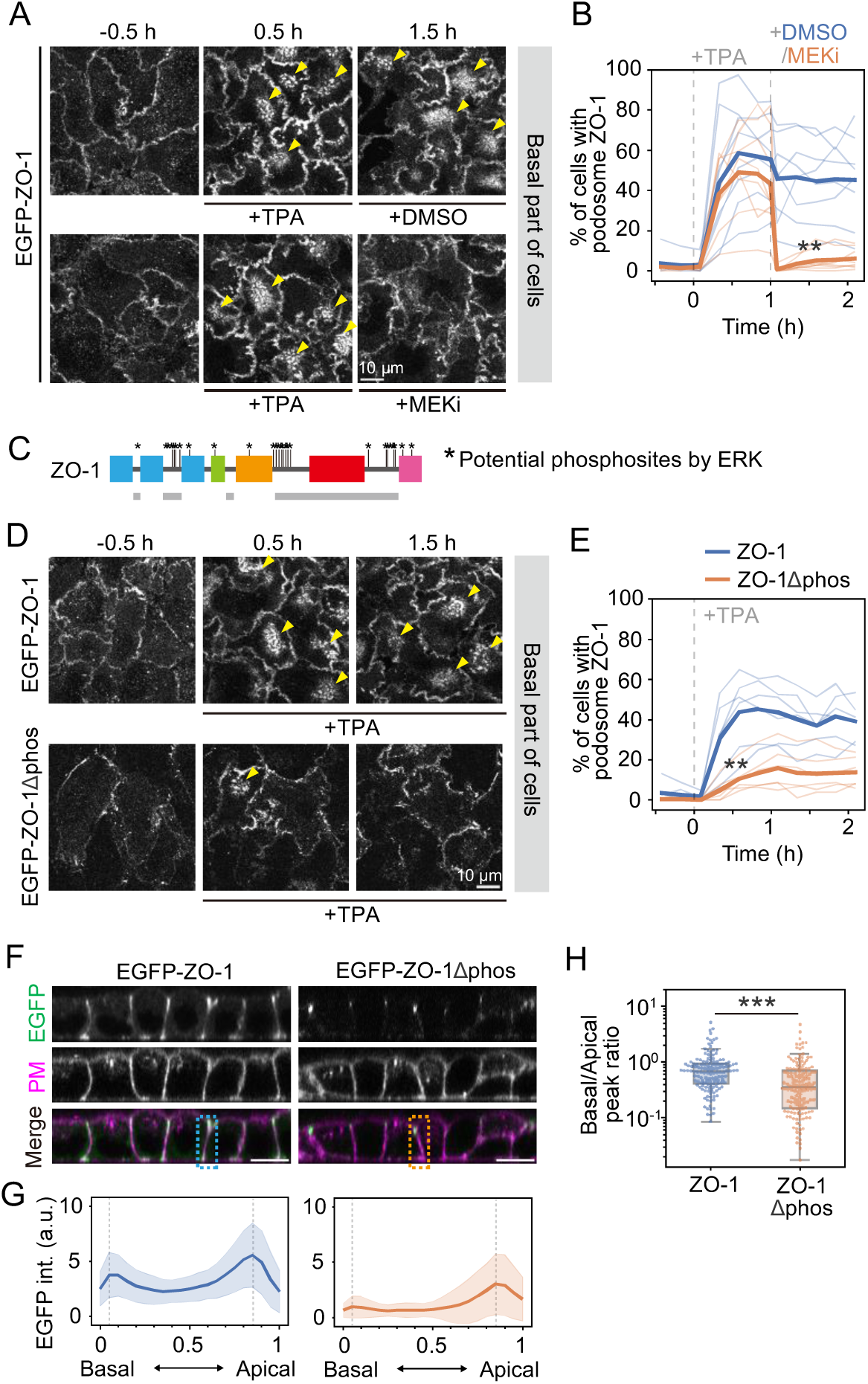
ERK phosphorylation- mediated ZO-1 translocation to podosomes (A) Time-lapse images of the basal part of the cells (MDCK II ZO-1/2 dKO/EGFP-ZO-1) treated with 10 nM TPA, followed by DMSO or 100 µM PD0325901 (a MEK inhibitor; MEKi). Arrowheads indicate clusters of podosomes. (B) Quantified data of panel A. Data are from four independent experiments. Thin lines and bold lines indicate the results of each sample (8 for each condition) and their average, respectively. (C) A schematic image of ZO-1 with the potential phosphorylation sites by ERK. (D) Time-lapse images of the basal part of the MDCK II ZO-1/2 dKO/EGFP-ZO-1 (upper) or MDCK II ZO-1/2 dKO/EGFP-ZO-1Δphos (lower) cells treated with 10 nM TPA. Arrowheads indicate clusters of podosomes. (E) Quantified data of panel D. Data are from three independent experiments. Thin lines and bold lines indicate the results of each sample (6 for each condition) and their average, respectively. (F) Lateral views of EGFP-ZO-1 and EGFP-ZO-1Δphos cell lines. PM, plasma membrane. (G) Quantified EGFP intensity along the lateral membranes. Data are from two independent experiments. The numbers of cell–cell interfaces were n = 141 (EGFP-ZO-1) and n = 145 (EGFP-ZO- 1Δphos). Lines and bands indicate the average and s.d., respectively. (H) The ratio of EGFP intensity at the basal to apical peak positions. The peak positions are indicated in panel G. Statistical significance was tested by the Mann-Whitney U-test.

To address the possibility that direct phosphorylation by ERK regulates the translocation of ZO- 1 to podosomes, we constructed a non-phosphorylatable mutant of ZO-1 by replacing all 26 potential ERK phosphorylation sites (*i.e.*, serine or threonine followed by proline; Fig. 4C) with alanine (ZO- 1Δphos). In a confluent monolayer cultured overnight, the EGFP-ZO-1Δphos was localized to TJs at a level comparable to EGFP-ZO-1 (Fig. S4A), suggesting that the introduced mutations did not cause loss of function of ZO-1 in forming TJs. When cells expressing either EGFP-ZO-1 or EGFP-ZO- 1Δphos were treated with TPA to induce podosome formation, EGFP-ZO-1 was accumulated in a large cluster of podosomes, while EGFP-ZO-1Δphos was localized to a small podosome cluster (Fig. 4D; Video S6). The percentage of cells with podosome ZO-1 was significantly lower in ZO-1Δphos- expressing cells than ZO-1-expressing cells (Fig. 4E). These results suggest that phosphorylation of ZO-1 promotes its localization to podosomes.

Next, we investigated the pathway through which ERK-phosphorylated ZO-1 translocates from TJs to podosomes. One day after seeding the cells, ZO-1 was widely distributed along the lateral membranes in the intercellular adhesions, with a predominant localization at the apical-most part of the cells (Fig. 4F, left). In contrast, ZO-1Δphos localization was restricted to the apical-most part (Fig. 4F, right). Quantification of fluorescence intensity along the lateral membranes revealed that EGFP-ZO-1 exhibited distinct peaks at both the apical and basal sides of the cells, while EGFP-ZO-1Δphos showed a distinct peak only at the apical side, with the basal peak being almost negligible (Fig. 4G). The ratio of fluorescence intensity at the basal to apical peak positions was lower for ZO-1Δphos than for ZO-1 (Fig. 4H). MEKi treatment also reduced the basal localization of ZO-1 compared to DMSO treatment (Fig. S4B-D). These results suggest that ERK-mediated phosphorylation promotes the movement of ZO-1 from the apical to the basal side of cells along the lateral membranes. We note that western blotting showed a lower expression level of ZO-1Δphos protein than of the ZO-1 or ZO-1ΔABR proteins in the stable cell lines (Fig. S5A and S5B). A decrease in ZO-1 protein level was also observed in cells treated with MEKi, although the decrease was not significant (Fig. S5C-E), suggesting that phosphorylation may increase the stability of the ZO-1 protein. Therefore, the differences in ZO-1 distribution along the lateral membranes observed in Figure 3F-H and S4B-D could be partly attributable to differences in the ZO-1 protein level. Nevertheless, the disappearance of ZO-1 from podosomes upon MEKi treatment was a rapid response occurring immediately after addition of the inhibitor, indicating that the effect of protein degradation is likely negligible in this process.

### ZO-1 at TJs facilitates ERK activation propagation and collective cell migration through the actin-binding region

We next investigated the effect of intracellular translocation of ZO-1 on collective cell migration by means of ZO-1 or ZO-1 mutant rescue experiments. To quantify the movement of cells within a population, we applied optical flow analysis to DIC images. In the parental MDCK II cells, intercellular propagation of cell movement was frequently observed (Fig. 5A), consistent with the previous reports^7,11^. In contrast, the ZO-1/2 dKO cells completely lost such propagation and exhibited markedly reduced displacement of the leading edge of the cell population (Fig. 5B). By expressing EGFP-ZO-1, the cells restored both the intercellular propagation of cell movement and the displacement of the leading edge (Fig. 5C). As indicated by the slope of the waves in kymographs, the propagation speed of cell movement in ZO-1/2 dKO + EGFP-ZO-1 cells (1.1 ± 0.2 μm/min) was slower than that of the parental MDCK II cells (2.0 ± 0.3 μm/min), possibly due to the absence of ZO-2 and/or the fusion of EGFP to ZO-1. We also examined whether cell motility could be restored by expressing ZO-1 mutants that exhibited reduced localization to podosomes. In EGFP-ZO-1ΔABR- expressing cells, the displacement of the leading edge was not as fast as in EGFP-ZO-1-expressing cells, and the intercellular propagation of cell movement was not restored (Fig. 5D). On the other hand, EGFP-ZO-1Δphos-expressing cells restored both the intercellular propagation of cell movement and the displacement of the leading edge (Fig. 5E). The propagation speed of cell movement (1.2 ± 0.2 μm/min) was comparable to that of EGFP-ZO-1-expressing cells. These results indicate that ZO-1 facilitates collective cell migration and that its interaction with F-actin is involved in this process.

**Figure 5.**
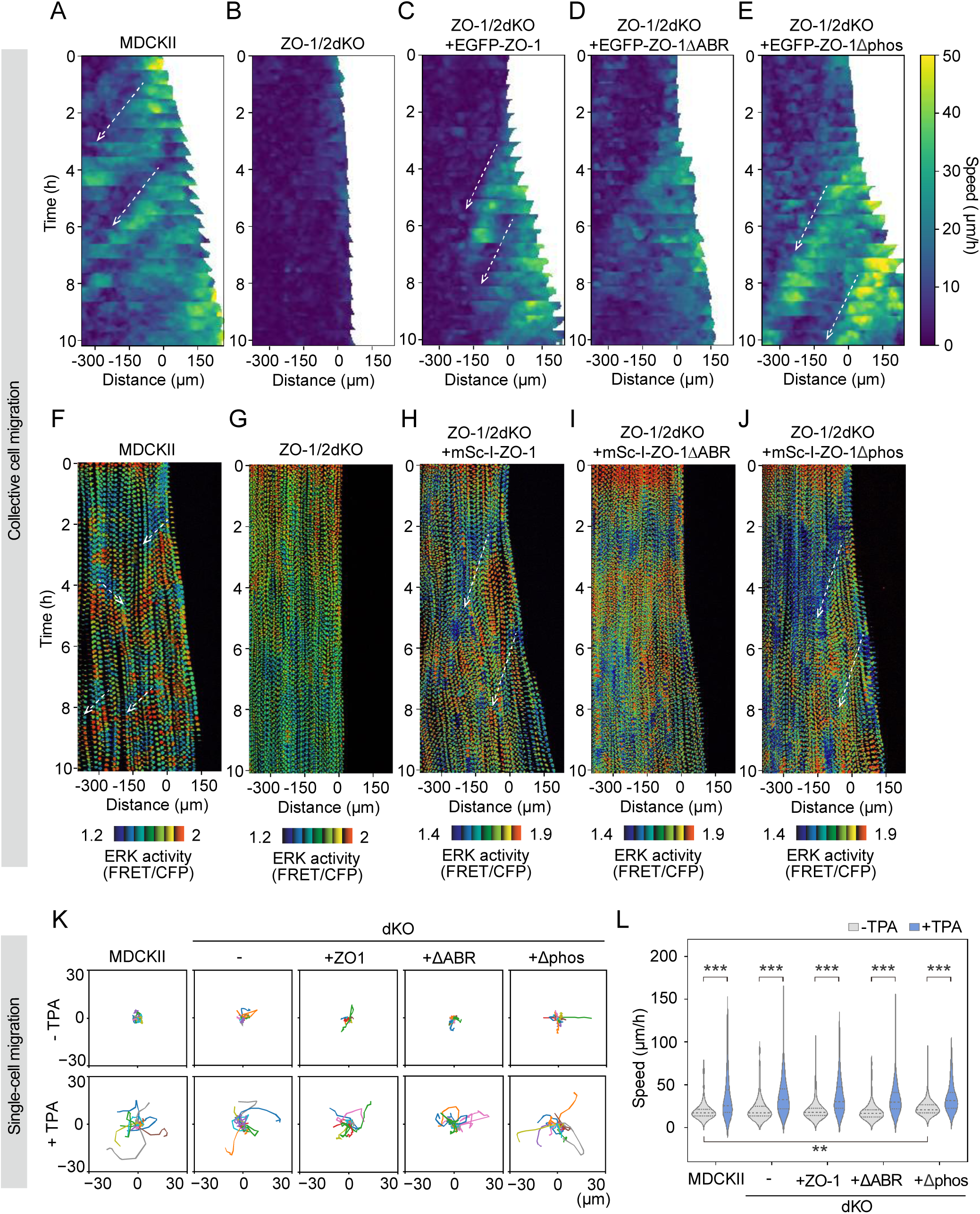
ERK activation propagation and collective cell migration facilitated by ZO-1 at TJs (A-E) Kymographs of the speed of cell movement quantified by optical flow analysis. Each graph shows representative data from three independent experiments for MDCK II (A), MDCK II ZO-1/2 dKO (B), MDCK II ZO-1/2 dKO/EGFP-ZO-1 (C), MDCK II ZO-1/2 dKO/EGFP-ZO-1ΔABR (D) or MDCK II ZO-1/2 dKO/EGFP-ZO-1Δphos (E), respectively. Dashed arrows indicate the propagation of collective cell migration. (F-J) Kymographs of ERK activity visualized by EKAREV-NLS. Each graph shows representative data from at least two independent experiments for MDCK II (F), MDCK II ZO-1/2 dKO (G), MDCK II ZO-1/2 dKO/mScarlet-I-ZO-1 (H), MDCK II ZO-1/2 dKO/mScarlet-I-ZO-1ΔABR (I) or MDCK II ZO-1/2 dKO/mScarlet-I-ZO-1Δphos (J), respectively. Dashed arrows indicate the propagation of ERK activation. (K) Trajectories of single-cell migration under the conditions with or without 10 nM TPA. Data of 20 representative cells from each sample are shown. (L) Single-cell migration speed under conditions with or without 10 nM TPA. n = 149 (MDCK II), n = 126 (dKO), n = 143 (dKO + ZO-1), n = 115 (dKO + ZO-1ΔABR), and n = 131 (dKO + ZO-1Δphos). The statistical significance of differences between the conditions with and without TPA for each cell line was tested by the Wilcoxon signed-rank sum test. The statistical significance of differences between cell lines under the conditions with or without TPA was tested by the Kruskal–Wallis test and the Steel–Dwass test.

Additionally, ZO-1 at TJs rather than at podosomes is likely important for facilitating collective cell migration because the expression of EGFP-ZO-1Δphos, which can not efficiently accumulate in podosomes, could restore the reduced collective migration of ZO-1/2 dKO cells.

To investigate the cause of the differences in intercellular propagation of cell movement, we examined ERK activation propagation. In collective migration of epithelial cells, the intercellular propagation of ERK activation propels the migration of the cell population in a direction opposite that of the ERK activation wave^7^. Based on this fact, we hypothesized that the absence of ZO-1 or the deletion of its ABR may disrupt the intercellular propagation of ERK activation, and thereby suppress the intercellular propagation of cell movement. To examine this possibility, we investigated ERK activity during collective cell migration using a FRET-based ERK activity sensor, as shown in Figure 3 (Video S7). In the parental MDCK II cells, intercellular propagation of ERK activation was observed throughout the cell population (Fig. 5F), and the propagation speed of ERK activation (2.5 ± 0.7 μm/min) was comparable to that of cell movement (2.0 ± 0.3 μm/min) and also to that in the previous report (2.12 μm/min)^7^. In ZO-1/2 dKO cells, the ERK propagation was completely lost (Fig. 5G). It was restored by expressing mScarlet-I-ZO-1 or mScarlet-I-ZO-1Δphos (Fig. 5H and 5J), although the propagation speed (ZO-1: 1.3 ± 0.5 μm/min; ZO-1Δphos: 1.3 ± 0.5 μm/min) was slower than that of the parental MDCK II cells (2.5 ± 0.7 μm/min). In contrast, expression of mScarlet-I-ZO-1ΔABR did not restore the ERK propagation (Fig. 5I). These results suggest that ZO-1 contributes to the intercellular propagation of ERK activation through its ABR, thereby promoting the intercellular propagation of cell movement during collective cell migration.

Next, to examine whether ZO-1 affects the motility of individual cells *per se*, we performed a single-cell migration assay, in which the MDCK II cells were sparsely seeded on a glass-based dish. Under the conditions without any drug treatment, there was no significant difference in migration speed between the parental MDCK II cells and the ZO-1/2 dKO cells (Fig. 5K and 5L). Therefore, the reduced migration speed of the ZO-1/2 dKO cells in collective migration was likely due to altered cell– cell interactions rather than to the motility of individual cells. In the cells expressing EGFP-ZO-1 or EGFP-ZO-1ΔABR in the ZO-1/2 dKO background, the single-cell migration speed was comparable to that of the parental MDCK II and ZO-1/2 dKO cells, while cells expressing EGFP-ZO-1Δphos exhibited a slightly higher migration speed (Fig. 5K and 5L). Although the reason for the increased migration speed in EGFP-ZO-1Δphos-expressing cells is unclear, this result suggests that ZO-1 may influence cell migration through mechanisms other than the formation of TJs or podosomes. To examine whether the localization of ZO-1 to podosomes affects the motility of individual cells, we investigated the single-cell migration speed under TPA-treated conditions. We found that the addition of TPA enhanced cell migration in all cell lines compared to the conditions without TPA, with no significant differences in migration speed between cell lines (Fig. 5K and 5L). This result suggests that the presence or absence of ZO-1 localization to podosomes does not affect the motility of individual cells, and that TPA enhances cell migration by a mechanism other than podosome formation.

### ZO-1 at podosomes contributes to traction force and cell invasion

To elucidate the function of ZO-1 localized at podosomes, we first focused on traction force. Podosomes consist of an actin-based core and a surrounding adhesive ring. As shown in Figure 1F, ZO- 1 is localized at the adhesive ring. Proteins constituting the adhesive ring, such as vinculin and paxillin, are known to contribute to traction force generation on the substrate^41^. Indeed, podosomes have been reported to generate traction forces ranging from 200 to 700 Pa depending on the substrate stiffness^42^. To investigate whether the ZO-1 localized at podosomes plays a role in generating traction forces, we quantified the traction force in EGFP-ZO-1- or EGFP-ZO-1ΔABR-expressing cells treated with TPA by traction force microscopy^43,44^. The cells were sparsely seeded on polyacrylamide gel containing fluorescent beads, and the traction force exerted by the cells was estimated based on the gel’s elasticity and the displacement of the fluorescent beads. Several tens of minutes after TPA treatment, approximately half of the cells showed podosomes, as shown in Figure 2E. In all cells, we observed the high traction forces in the periphery of cells, possibly due to the lamellipodial extension induced by TPA treatment (Fig. 6A). In particular, the regions with podosomes exhibited higher traction forces than other regions of the cells (Fig. 6A, arrowheads). EGFP-ZO-1ΔABR, which did not form the adhesive ring even after TPA treatment as shown in Figure 2, demonstrated low traction forces in the interior region of the basal cell surface (Fig. 6A). We compared the average traction force across the basal surface of cells, and found that EGFP-ZO-1-expressing cells with podosomes showed higher traction forces than those without podosomes (Fig. 6B). Furthermore, the traction force exerted by EGFP-ZO-1ΔABR-expressing cells was significantly lower than that of EGFP-ZO-1-expressing cells. Considering that EGFP-ZO-1ΔABR-expressing cells form the actin core but not the adhesive ring of podosomes (Fig. 2H), these results suggest that the adhesive ring, rather than the core, is crucial for generating traction forces, and that ZO-1 enhances traction forces by regulating the adhesive ring formation.

**Figure 6.**
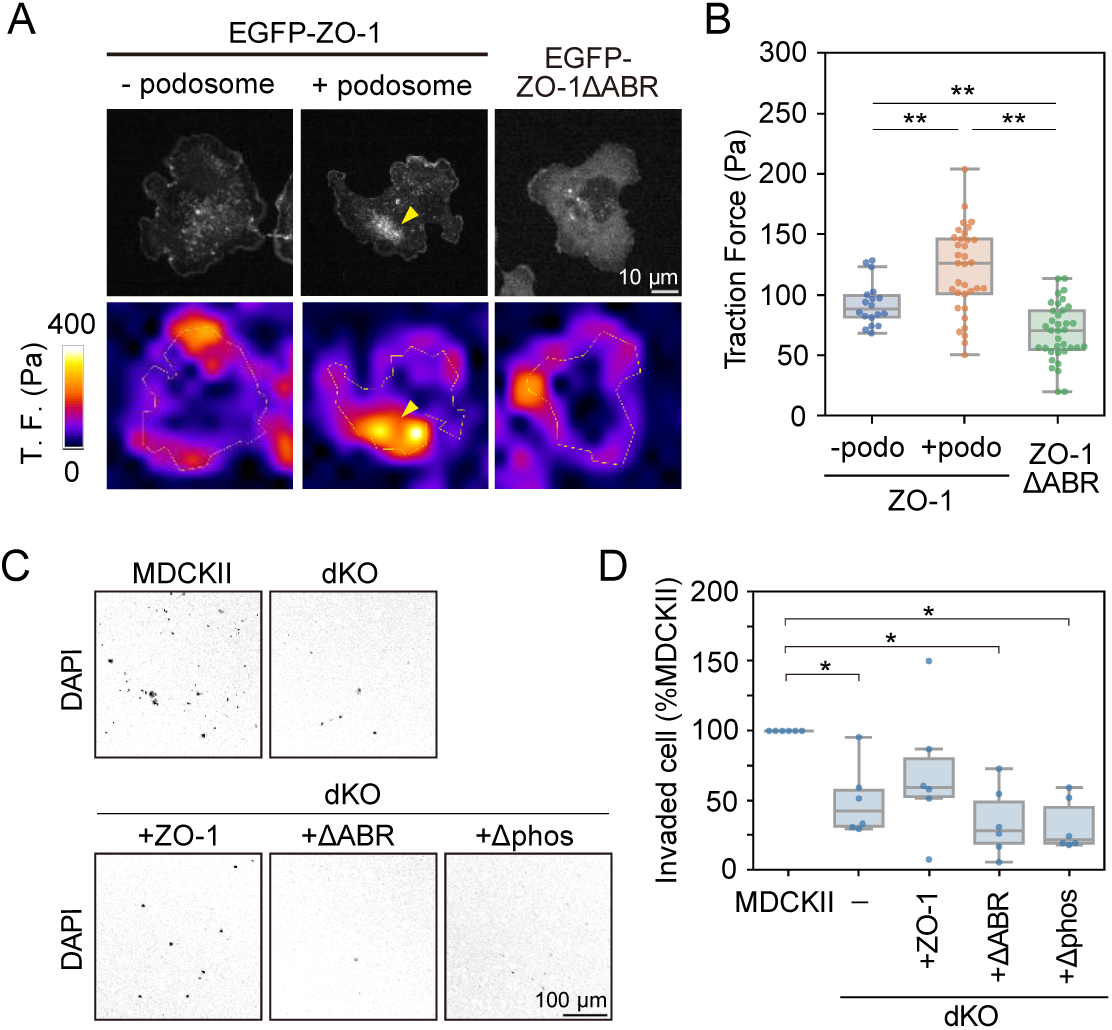
Contribution of ZO-1 to traction force and cell invasion (A) Fluorescence images of cells treated with TPA (upper panels) and traction forces estimated from the gel’s elasticity and the displacement of fluorescent beads embedded in the gel (lower panels). Arrowheads indicate the podosome region. (B) Quantified data of panel A. Each dot indicates the average traction force over the whole basal surface of individual cells. N = 18 (MDCK II ZO-1/2 dKO/EGFP-ZO-1 - podosome), n = 34 (MDCK II ZO-1/2 dKO/EGFP-ZO-1 + podosome), and n = 36 (MDCK II ZO-1/2 dKO/EGFP-ZO-1ΔABR) from three independent experiments. (C) Inverted fluorescence images of invaded cells on the lower surface of porous membranes. (D) The number of invaded cells normalized with that of the parental MDCK II cells. Each dot indicates an average data point from six independent experiments. Statistical significance was tested by the Kruskal-Wallis test and the Steel-Dwass test.

Finally, we investigated the contribution of ZO-1 to cell invasiveness. ECM degradation and promotion of cell invasion are fundamental functions of podosomes^20^. To examine whether ZO-1 localization to podosomes affects the invasiveness of cells, we performed a trans-well invasion assay. In this assay, TPA-treated cells were sparsely seeded on a porous membrane coated with Matrigel. FBS was added to the medium below the membrane as an attractant, inducing invasive cells to migrate through the membrane pores to the underside of the membrane by degrading the Matrigel. To evaluate the invasiveness of cells, we counted the number of cells that had moved to the lower surface of the membrane. We found that there was a significantly lower number of invading ZO-1/2 dKO cells compared to parental MDCK II cells (Fig. 6C), with the invaded cell count of ZO-1/2 dKO cells being less than half that of the parental cells (Fig. 6D). Expression of EGFP-ZO-1 partially restored the invasiveness of the dKO cells, to 60–70% of the level in parental cells. In contrast, the expression of mutant ZO-1 proteins (EGFP-ZO-1ΔABR or EGFP-ZO-1Δphos), which poorly localize to podosomes, failed to restore the invasiveness. These results indicate the importance of ZO-1 localization to podosomes for promoting cell invasion.

## Discussion

In this study, we revealed that ZO-1 dynamically changes its localization from TJs to podosomes during collective cell migration and plays a distinct role from its well-known function in TJ formation. In collectively migrating epithelial cells, ZO-1 transiently dissociates from TJs and accumulates at the basal surface of the cells, contributing to the formation of circular assemblies or irregular clusters of podosomes (Fig. 7A, green arrows). The translocation of ZO-1 from TJs to podosomes is driven by ERK activation, probably mediated by direct phosphorylation of ZO-1 by ERK (Fig. 7A, orange arrows). Phosphorylated ZO-1 localizes to the adhesive ring of podosomes by interacting with F-actin (Fig. 7B). ZO-1 localized at the adhesive ring then enhances traction force generation and cell invasion by contributing to the ring formation. Additionally, ZO-1 plays not only a role downstream of ERK but also a role in propagating ERK activation between cells in a migrating cell population. This process involves ZO-1 at TJs, rather than at podosomes, where it mediates intercellular propagation of ERK activation (Fig. 7A, dotted orange arrows) and consequent cell movement via a mechanism involving its ABR.

**Figure 7.**
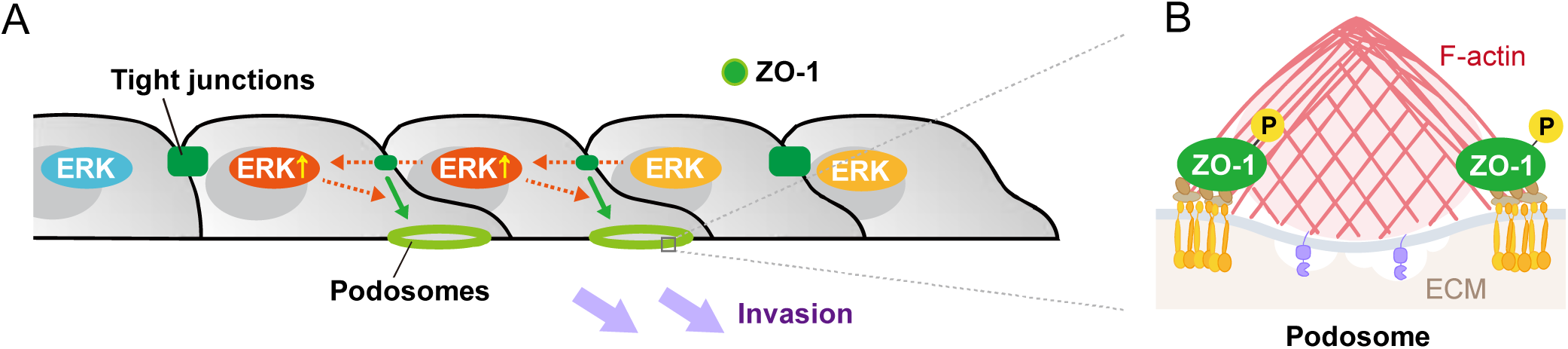
ZO-1 translocation from TJs to podosomes (A) A schematic image of ZO-1 translocation from TJs to podosomes during collective cell migration. (B) A schematic image of a podosome with ZO-1 at the adhesive ring.

In intact epithelial cell sheets, ZO-1 forms TJs at the apical side of the cell–cell interface. In this study, we demonstrated that ZO-1 dynamically dissociates from TJs and translocates to podosomes during collective cell migration. Dissociation of ZO-1 from TJs has also been reported during epithelial–mesenchymal transition (EMT)^45,46^. In addition, a previous study reported that expression of a truncated mutant of ZO-1 that does not localize to TJs can induce typical EMT changes, such as reduced cell–cell adhesion factors and increased vimentin^47^. Our work demonstrated that ERK activity regulates ZO-1 translocation from TJs to podosomes. A non-phosphorylatable mutant of ZO-1 was confined to TJs and did not spread to other cellular regions (Fig. 4F-H), suggesting that ERK-mediated phosphorylation promotes ZO-1 dissociation from TJs. Honigmann and his colleagues recently demonstrated that TJ formation is driven by liquid–liquid phase separation (LLPS) of ZO-1 on plasma membranes and that lower phosphorylation levels facilitate LLPS and therefore the accumulation of ZO-1 to TJs^48^. They explored ZO-1 phosphorylation by casein kinase II (CK2) and identified 47 phosphorylation sites. Of these, 11 sites overlap with the potential ERK phosphorylation sites that were a focus in this study. Furthermore, most of these sites are located in the intrinsically disordered regions (IDRs), which generally play important roles in liquid-liquid phase separation (LLPS) of biomolecules^49^. Phosphorylations of IDRs promote or suppress LLPS by affecting electrostatic interactions between biomolecules, depending on the specific proteins and contexts^50^. ERK-mediated phosphorylation of ZO-1 may modulate the charge distribution of the IDRs, thereby dissolving LLPS and promoting ZO-1 dissociation from TJs.

We demonstrated that ZO-1 accumulates at the adhesive ring of podosomes via its actin-binding region. While we used epithelial cells, a previous study reported ZO-1 localization in the adhesive ring of immune-cell podosomes^51^, which are commonly used in podosome studies. The presence of actin cores of podosomes in cells lacking ZO-1 localization to podosomes (Fig. 2H) suggests that ZO-1 is not essential for core formation. Instead, our results indicated that ZO-1 plays a role in ring formation, based on the fact that the loss of ZO-1 in the adhesive ring suppressed the localization of another ring-forming protein, vinculin (Fig. S3). Previous studies have reported that ZO-1 interacts with integrins^52,53^ and vinculin^54^, both of which are adhesion-related proteins and known to be components of the adhesive ring of podosomes^55,56^. Combining our results with the previously reported findings, we propose that ZO-1 functions to link integrins and other ring-forming proteins at podosomes, similar to its role in bridging claudins and cytoplasmic proteins at TJs. Moreover, we hypothesize that ZO-1 is involved in the mechanoresponsive functions of podosomes that facilitate cell invasion. In the ring of podosomes, adhesion complexes connect actomyosin bundles to the substrate, transmitting the contractile force of actomyosin to the substrate and generating the pulling force^20^. On the other hand, in the core of podosomes, matrix metalloproteinases degrade the ECM, while the polymerization of branched actin filaments generates protruding forces against the substrate^20^. Simulations have suggested that the pulling force in the ring is important for the formation of protrusions by podosomes^25,26^, and indeed, inhibition of myosin activity has been shown to reduce protruding forces and ECM degradation^23,57,58^. We here demonstrated that the absence of ZO-1 from the ring of podosomes reduced cell invasion (Fig. 6C and 6D), suggesting that the loss of the linkage between integrins and other ring-forming proteins disrupted the force transmission through the adhesive ring and thereby the formation of protrusions by podosomes.

While ZO-1 at podosomes promotes cell invasion, ZO-1 at TJs facilitates collective cell migration (Fig. 5). We previously demonstrated that intercellular waves of ERK activation drive collective cell migration in the direction opposite to the wave itself^7^. Our present rescue experiments suggest that ZO-1 promotes migration by participating in the intercellular propagation of ERK activation (Fig. 5). Further, ZO-1ΔABR expression did not completely restore intercellular waves of ERK activation and collective cell movement (Fig. 5). These results indicate that ZO-1 mediates ERK activation propagation through its interaction with the actin cytoskeleton. Previous studies have demonstrated that ZO-1 undergoes conformational changes under the tensile stress induced by actomyosin^59,60^. The regions exposed by the conformational changes contain the vinculin-binding site^54^, suggesting that ZO-1 acts as a tension-transducer. Similarly, α-catenin, a component of adherens junctions, changes its conformation in a tension-dependent manner and transmits tensile force between cells^61^. The knockout of α-catenin disrupts organized ERK propagation during collective cell migration^11^. Taking all these findings into account, ZO-1 likely acts as a tension transducer between cells to propagate ERK activation and facilitate collective cell migration. In addition, we note that ZO- 1 contributes to tissue fluidity during collective cell migration. Previous studies have indicated that the depletion of ZO proteins results in hyperdevelopment of actomyosin bundles at the cell periphery and that ZO proteins are necessary for maintaining tissue fluidity^14,62^. The recovery of collective cell migration speed observed with the expression of ZO-1 or its mutants can be attributed, at least in part, to the restoration of tissue fluidity.

This study provides new insights into the dynamic behavior of ZO-1 and its role in cell adhesion and movement. Despite these advancements, several significant points remain to be elucidated. First, the pathway of ZO-1 translocation from TJs to podosomes is unclear. Although we propose that ZO-1 spreads to the basal side of the cells along the lateral membranes upon phosphorylation, ZO-1 protein stability seems to be also affected by phosphorylation (Fig. S5). We have previously demonstrated that ZO-1 forms cytoplasmic condensates, a structure distinct from both TJs and podosomes^34^. The formation of the condensates is thought to be driven by LLPS of ZO-1. The formation, dissolution, and/or intracellular movement of these condensates can be considered as potential mechanisms regulating ZO-1 translocation from TJs to podosomes. Secondly, while we focused on ERK activity in this study, the regulation of ZO-1 localization by other kinases should also be taken into account. The amino acid sequences targeted for phosphorylation by ERK (*i.e.*, serine or threonine followed by proline) might be phosphorylated by other MAP kinases such as p38 and JNK. These kinases are known to be activated by inflammatory cytokines and various cellular stresses^63^, suggesting that the localization changes of ZO-1 revealed in this study are involved in the cellular responses to these stimuli. By addressing these questions we can achieve a profound understanding of the multifaceted roles of ZO-1 in regulating cellular functions *in vivo*.

## Supporting information

Key resources table

## Acknowledgements

We thank the Exploratory Research Center on Life and Living Systems (ExCELLS) for the imaging support with the ZEISS Lattice Lightsheet 7. This study was supported by the Japan Society for the Promotion of Science (JSPS) (grant number 23KJ2217 to S.H.; no. 21H02523 to M.F.; no. 21H02493 to N.U.; nos. JP22H02625 and JP24H01416 to K.A.), and by grants from the International Research Collaboration Center at the National Institutes of Natural Sciences (to S.H., N.U., and K.A.). T.O. and K.A. were supported by Takeda Science Foundation.

## Author contributions

Conceptualization, S.H., N.U., K.A.; Investigation, S.H., Y.K.; Writing - Original Draft, S.H.; Writing - Review & Editing, S.H., Y.K., N.K., T.O., M.F., N.U., K.A.; Visualization, S.H., Y.K.; Project Administration, N.U., K.A.; Funding Acquisition, S.H., N.U., K.A.

## STAR methods

### Resource availability

#### Lead contact

Further information and requests for resources and reagents should be directed to and will be fulfilled by the lead contact, Kazuhiro Aoki.

#### Materials availability

Plasmids and cell lines generated in this study are available upon request.

#### Data and code availability

All original data and code are available upon request.

### Experimental model and study participant details

#### Cell culture

MDCK II cells were maintained in Dulbecco’s modified Eagle’s medium (DMEM, high glucose; Nacalai Tesque, 08459-64) supplemented with 10% fetal bovine serum (FBS; Sigma-Aldrich, F7524), in a 5% CO2 humidified incubator at 37°C. At passage, the growth medium was removed and 8–10 mL of 1 mM EDTA/PBS was added. After 10 min incubation at 37°C, the 1 mM EDTA/PBS was removed and 1 mL of 0.25% trypsin/0.02% EDTA/PBS was added. The trypsin/EDTA solution was removed immediately and the cells were incubated for 10 min at 37°C. The cells were resuspended in the pre-warmed growth medium.

#### Establishment of stable cell lines

A lentiviral system was used to establish stable cell lines expressing EGFP-ZO-1, EGFP-ZO-1ΔABR, EGFP-ZO-1Δphos, H2B-iRFP or mScarlet-I-KRasCT. To prepare the lentivirus, the pCSII-based lentiviral vector^64^, psPAX2 (Addgene plasmid #12260), and pCMV-VSV-G-RSV-Rev were cotransfected into Lenti-X 293T cells. MDCK II cells were cultured in the culture medium containing the lentivirus and 10 µg/mL polybrene (Nacalai Tesque) for 3–6 h. Instead of lentiviral transfer, a PiggyBac or Tol2 transposon system was also used to establish stable cell lines expressing EKAREV-NLS, mScarlet-I-ZO-1, mScarlet-I-ZO-1ΔABR, or mScarlet-I-ZO-1Δphos. The pPBbsr2- and pT2A- based vector were cotransfected with pCAGGS-hyPBase and pCAGGS-T2TP encoding the Piggybac and Tol2 transposase, respectively, by using an Amaxa nucleofector system (Lonza) at a 4:1 ratio. Cells were selected by using the following antibiotics: puromycin, 2 µg/mL; zeocin, 100 µg/mL; hygromycin, 100 µg/mL; blasticidin S, 5∼10 µg/mL. Single-cell clones were further isolated by limited dilution.

### Method details

#### Plasmid construction

All plasmids constructed and used in this study are summarized in the Key Resources table. The cDNA of human ZO-1 and ZO-1ΔABR were amplified by PCR, and subcloned into a pCSIIpuro vector carrying a puromycin-resistance gene to produce pCSIIpuro-EGFP-ZO-1 and pCSIIpuro-EGFP-ZO- 1ΔABR, respectively. To generate the non-phosphorylatable mutant (pCSIIpuro-EGFP-ZO-1Δphos), all serines and threonines in the serine-proline (SP) and threonine-proline (TP) sequence in ZO-1 were replaced with alanine. Three regions with many mutation sites (a.a. 274–414, a.a. 857–986, and a.a. 1410–1579) were synthesized *de novo* by FASMAC, and all the other mutations were introduced by PCR. The mutated sites (S168, S315, S329, T354, T363, T379, S402, S454, S531, T709, T868, S899, S912, S927, S933, T960, S968, T976, T1425, T1521, T1528, S1545, T1567, S1570, S1617, and S1695) are annotated in the sequence of the Benchling link. ZO-1, ZO-1ΔABR, and ZO-1Δphos were subcloned into the pT2Apuro vector fused with mScarlet-I to generate pT2Apuro-mScarlet-I-ZO-1, pT2Apuro-mScarlet-I-ZO-1ΔABR, and pT2Apuro-mScarlet-I-ZO-1Δphos, respectively. psPAX2 was kindly provided by Didier Trono (Addgene plasmid # 12260). pCMV-VSV-G-RSV-Rev was kindly provided by Hiroyuki Miyoshi^64^. pCAGGS-T2TP was kindly provided by Koichi Kawakami. KRas C-terminus (KRasCT)^10^ and mScarlet-I were subcloned into a pCSIIhyg vector harboring a hygromycin-resistance gene to produce pCSIIhyg-mScarlet-I-KRasCT.

#### Fluorescence imaging

Cells were imaged using an IX83 inverted microscope (EVIDENT) equipped with an FV3000 confocal unit (EVIDENT) and diode lasers. An oil immersion objective lens (UPlanXApo 100XO, N.A. 1.45; UPLlanXApo 60XO, N.A. 1.42; EVIDENT) and an air/dry objective lens (UPlanXApo 20X, N.A. 0.8; UPlanSApo 10X, N.A. 0.4; EVIDENT) were used. The excitation laser and fluorescence filter settings were as follows: excitation laser, 405 nm (Alexa Fluor 405, DAPI), 488 nm (EGFP, Alexa Fluor 488), 561 nm (Alexa Fluor 555, iFluor 555, CellMask Orange), and 640 nm (iRFP670, Alexa Fluor 633); dichroic mirror, DM 405/488/561/640; emission filters, SDM400-470 (Alexa Fluor 405, DAPI), SDM400-540 (EGFP, Alexa Fluor 488), SDM400-580 (Alexa Fluor 555, iFluor 555, CellMask Orange), and SDM400-620 (iRFP670, Alexa Fluor 633). The microscope was controlled by FLUOVIEW31S-SW software (ver.2.6.1.243). During observation, cells were incubated in a stage incubator set at 37°C and 5% CO2 (STXG-IX3WX, Tokai Hit).

For the single-cell migration and FRET imaging shown in Figure 5, cells were imaged using an IX83 inverted microscope (EVIDENT) equipped with an sCMOS camera (Prime: Photometrics, Tucson, AZ) and SPECTRA X Light Engine (Lumencor). An air/dry objective lens (UPLXAPO 20X, N.A. 0.8; UPLXAPO 10X, N.A. 0.4; EVIDENT) was used. The filter settings were as follows: excitation filter, 438/24 nm (CFP, FRET) and 632/22 nm (iRFP); dichroic mirror, FF458 (CFP, FRET) and FF409/493/573/652/759 (iRFP); emission filter, FF01-483/32-25 (CFP), FF02-520/28 (FRET), and BLP01-664R (iRFP). The microscope was controlled by MetaMorph software (ver. 7.10.4.407).

During observation, cells were incubated in a stage incubator set to 37°C and 5% CO2 (STXG-IX3WX, Tokai Hit).

For the traction force microscopy and FRET imaging shown in Figure 3, cells were imaged using an IX83 inverted microscope (EVIDENT) equipped with an sCMOS camera (ORCA-Fusion BT: Hamamatsu Photonics), a spinning disk confocal unit (CSU-W1; Yokogawa Electric Corporation), and diode lasers. An oil immersion objective lens (UPLXAPO60XO, N.A. 1.42; EVIDENT) and an air/dry objective lens (UPLXAPO 20X, N.A. 0.8; EVIDENT) were used. The excitation laser and fluorescence filter settings were as follows: excitation laser, 445 nm (CFP, FRET), 488 nm (EGFP), and 561 nm (mScarlet-I, red fluorescent beads); dichroic mirror, DM 445/514/640 (CFP, FRET) and DM 405/488/561/640 (EGFP, mScarlet-I, red fluorescent beads); emission filters, 482/35 (CFP), 525/50 (FRET, EGFP), and 617/73 (mScarlet-I, red fluorescent beads). The microscope was controlled by MetaMorph software (ver. 7.10.3). During observation, cells were incubated in a stage incubator set at 37°C and 5% CO2 (STXG-IX3WX, Tokai Hit).

#### Confinement release assay

MDCK II cells (70 μL of 7x10^5^ cells/mL) were seeded in a 2-well culture insert (ibidi, 80209) on a 35 mm glass-based dish (IWAKI, 3910-035). After 24 h of incubation, the insert was removed. Cells were imaged using an FV3000 confocal microscope (EVIDENT). Images were displayed as a single z-slice for the basal side of the cells (Fig. 1A) and as a maximum intensity projection of multiple images for the apical side (Fig. 1B).

#### Immunostaining

Cells were fixed with 3.7% formaldehyde in PBS for 15 min at room temperature. After one wash with 0.05% Tween-20 in PBS, the cells were permeabilized with 0.2% Triton X-100 in PBS for 15 min at room temperature. After three washes with 0.05% Tween-20 in PBS, the cells were incubated in the blocking buffer (0.02% Triton X-100/3% BSA in PBS) for 1 h at room temperature. The cells were then incubated with primary antibodies in the blocking buffer overnight at 4 °C. The primary antibodies and the dilution rates were as follows: anti-ZO-1 (Invitrogen, 33-9100; 1/100), anti-TKS5 (Sigma-Aldrich, 09-268; 1/1000), and anti-vinculin (Proteintech, 26520-1-AP; 1/500). After three washes with 0.05% Tween-20 in PBS, the cells were incubated with secondary antibodies in the blocking buffer for 1 h at room temperature. The secondary antibodies and the dilution rate were as follows: Alexa Fluor 488-conjugated anti-mouse IgG (Invitrogen, A32723; 1/1000), Alexa Fluor 405-conjugated anti-rabbit IgG (Invitrogen, A48254; 1/1000), and Alexa Fluor 555-conjugated anti-rabbit IgG (Invitrogen, A32732; 1/1000). The cells were washed with 0.05% Tween-20 in PBS three times. For staining of actin filament, the cells were incubated with Alexa Fluor 633-conjugated phalloidin in PBS (Invitrogen, A22284; 1/4000) for 20 min or iFluor 555-conjugated phalloidin in 1% BSA in PBS (AAT Bioquest, 23119; 1/1000). The cells were washed with PBS three times and imaged with an FV3000 confocal microscope (EVIDENT).

#### Drug treatment

Cells were seeded on a 4-well glass-based dish (Greiner Bio-One). After overnight incubation, the cells were treated with 10 nM TPA or 100 µM PD0325901 to investigate the effect of PKC activation or MEK inhibition, respectively. Mock experiments were performed with DMSO treatment. Time-lapse images were taken using an FV3000 confocal microscope (EVIDENT). Nuclei were detected from H2B-iRFP fluorescence images using a Fiji plugin, StarDist^65^. The number of podosome clusters was counted by visual inspection. The percentage of the cells with podosomes was calculated by dividing the number of podosome clusters in the field of view by the number of nuclei in the same field of view.

#### Traction force microscopy

The gel solution was prepared with 10% acrylamide, 0.25% bisacrylamide, 0.8% ammonium persulfate, 0.08% TEMED (Nacalai Tesque), and 0.1% red fluorescent carboxylate-modified beads (0.2 μm diameter; Thermo Fisher Scientific, F8810). 13 μL of the mixture was placed on a 35 mm glass-based dish (IWAKI, glass 27 mm ø) and covered with a glass coverslip (Matsunami, 15 mm ø). After polymerization for 1 h at room temperature, the coverslip was removed, and 200 μL of 4 mM sulphosuccinimidyl-6-(4-azido-2-nitrophenylamino) hexanoate (Sulfo-SANPAH; Pierce) in 0.1 M HEPES buffer was added to the dish. The dish was exposed to UV light for 10 min, washed once with 0.1 M HEPES buffer, washed once with PBS, and then incubated in 0.1 mg/mL type-I collagen solution (Nitta Gelatin) at 4°C overnight. The collagen-coated dish was washed twice with PBS, and soaked in the culture medium for at least 6 h before being seeded with the cells. 2x10^4^ cells were seeded on the dish and incubated overnight before being treated with 10 nM TPA. Several tens of minutes after TPA treatment, the cells and the fluorescent beads were imaged using a spinning disk confocal microscope. The cells were then removed with 0.25% trypsin/0.02% EDTA/PBS. After acquiring images of only the beads, the displacement of the beads in the presence of cells when compared to the absence of cells was analyzed using Fiji plugins: Template Matching and iterative PIV. The traction force was estimated from the beads displacement and the gel elasticity using a Fiji plugin, FTTC. Note that the Young’s modulus of the gel was estimated to be ∼2 kPa according to a previous report^66^. The ROI was determined manually as the whole cell region where the EGFP signal was observed, and the average traction force in the ROI was recognized as the traction force of each cell.

#### Protein distribution along the lateral plasma membrane

To detect the plasma membrane, the cells were stained with CellMask Orange Plasma Membrane Stain (Invitrogen) according to the manufacturer’s protocol. In brief, the cells were washed once with PBS, and incubated in 1/1000 CellMask Orange solution in the culture medium for 10 min at 37°C. The cells were washed three times with PBS, and the imaging medium was added: 5% FBS (Sigma-Aldrich, F7524)/1xGlutaMAX (Gibco, 35050-061) in FluoroBrite DMEM (Thermo Fisher Scientific, A1896701). Cells were imaged using an FV3000 confocal microscope (EVIDENT). The line ROIs along the lateral membranes were manually determined from the CellMask Orange images, and the same ROIs were used to quantify EGFP distribution.

#### Single-cell migration

5x10^3^ cells were seeded in each well of a 4-well glass-based dish (Greiner Bio-One). After overnight incubation, the cells were imaged using an IX83 inverted microscope (EVIDENT) with a time interval of 5 min. One hour after the start of the imaging, a TPA solution was added to achieve a final concentration of 10 nM. The cells that remained in a single-cell state throughout the imaging were tracked using a Fiji pugin, LIM Tracker^67^.

#### Optical flow analysis

DIC images of collectively migrating cells were obtained using an FV3000 confocal microscope (EVIDENT). Optical flow was analyzed using Dense Optical Flow in OpenCV^68^ in Python. Mask images were generated using Fiji with the following procedures: After converting the original image to an 8-bit image, the Fiji functions “Find Edges” and “Gaussian Blur” were applied. The image was then binarized and filled with small empty areas using “Analyze Particle”.

#### Trans-well Invasion assay

A Matrigel-coated invasion chamber (Corning, 354480) and a 24-well companion plate (Corning, 353504) were rehydrated with serum-free DMEM (Nacalai Tesque, 08459-64) for at least 2 h at 37°C before seeding cells. Cells were suspended in serum-free DMEM at a concentration of 5x10^4^ cells/mL. TPA solution was added to the cell suspension to achieve a final concentration of 10 nM. After removal of serum-free medium, 500 μL of the cell suspension was added to the invasion chamber. The chamber was then set on a 24-well companion plate containing 750 μL of 10% FBS (Sigma-Aldrich, F7524)/DMEM. After 72 h incubation at 37°C with 5% CO2, non-invaded cells on the upper surface of the invasion chamber were removed by scrubbing with cotton swabs. The cells on the lower surface of the invasion chamber were washed once with PBS and then fixed with 3.7% formaldehyde in PBS for 15 min at room temperature. After being washed with PBS, the cells were incubated in 1 μg/mL DAPI in PBS for 20 min at room temperature. After a final wash with PBS, the cells were soaked with 50% glycerol in PBS for 5 min and imaged using an FV3000 confocal microscope (EVIDENT). Nuclei were detected using a Fiji plugin, StarDist^65^.

#### Western blotting

Cells were lysed in 1x SDS sample buffer. After sonication and boiling, the samples were separated by 5%–20% gradient SDS-polyacrylamide gel electrophoresis (Nacalai Tesque) and transferred to polyvinylidene difluoride membranes (Millipore). After blocking with Odyssey Blocking Buffer-TBS (LICOR Biosciences) for 1 h, the membrane was incubated with primary antibodies overnight at 4°C. After being washed with TBST three times, the membrane was incubated with secondary antibodies for 1 h at room temperature. After three washes with TBST, the membrane was imaged with an Odyssey infrared scanner (LI-COR Bioscience). For primary antibodies, anti-ZO-1 (Invitrogen, 33-9100; 1/1000) and anti-α-Tubulin (MBL, PM054; 1/1000) were diluted in Odyssey Blocking Buffer-TBS. For secondary antibodies, IRDye680RD-conjugated goat polyclonal anti-rabbit IgG (H + L) (1/5000; LI-COR Bioscience, 926-68071) and IRDye800CW-conjugated donkey polyclonal anti-mouse IgG (H + L) (1/5000; LI-COR Bioscience, 925-32212) were diluted in Odyssey Blocking Buffer-TBS.

#### Automatic segmentation of the circular assemblies of podosomes

We used U-Net^69^, a deep-learning model for semantic segmentation of biomedical images, for segmentation of the circular assemblies of podosomes in fluorescence images of mScarlet-I-ZO-1. For preparing training data, we manually annotated circular assemblies of podosomes in 20 images with 512–512 pixels (Fig. S6A). 14 out of 20 annotated images were used for training, 3 images were for validation, and 3 images were for testing. Upon training, three successive frames of fluorescence images were given to U-Net in order to predict a segmented image for the middle time point. We adopted a PyTorch implementation of U-Net (https://github.com/milesial/Pytorch-UNet), and used the Adam optimizer (batch size=8, learning rate=0.0003, 160 epochs) to minimize the binary cross-entropy loss. The procedure for data augmentation was implemented by using the Albumentations library (version 1.4.4) (See Fig. S6B and S6C for the learning curve and the settings of data augmentation).

The best weights with the lowest validation loss achieved a Dice score of 0.66 on the test images and were used for further analysis.

### Quantification and statistical analysis

#### Image analysis

For the quantification of the fluorescence images, the background was subtracted using the rolling-ball method adopted in Fiji. The efficiency of FRET was quantified as the ratio of fluorescence intensity of FRET to that of CFP. The FRET/CFP ratio images were visualized by using the intensity-modulated display (IMD) mode, where eight colors from red to blue are used to represent the FRET/CFP ratio, with the intensity of each color indicating the intensity of CFP.

#### Statistical analysis

Statistical analyses were performed with Python. The sample sizes, statistical tests, and *p*-values are indicated in the figures and the figure legends. p-values were classified as follows: ∗*p* < 0.05, ∗∗*p* < 0.01, ∗∗∗*p* < 0.001, and n.s. (not significant, i.e., *p* > 0.05).

