## Supplementary Information for "ZO-1 shuttles between tight junctions and podosomes by riding ERK activation waves during collective cell migration"

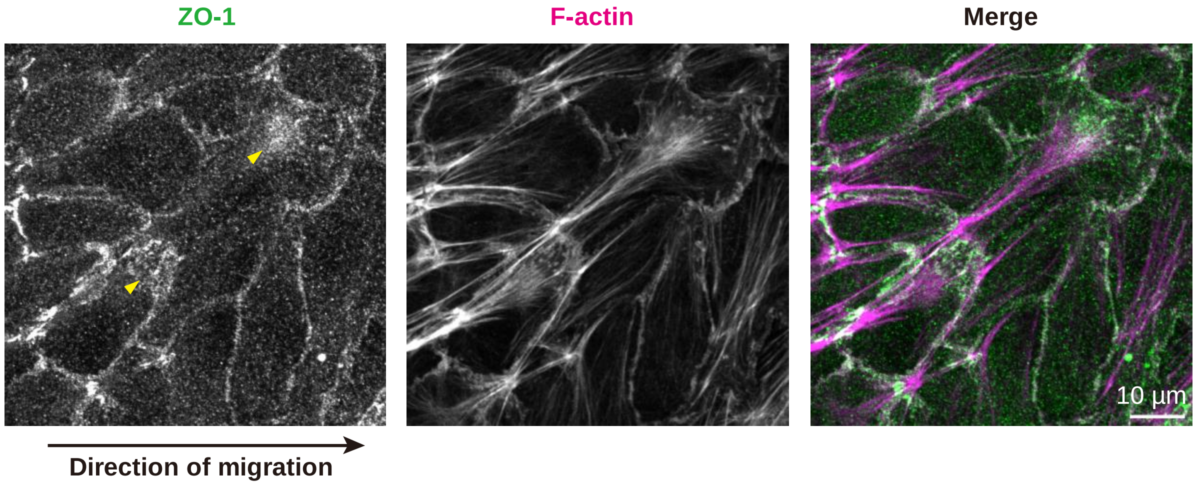


**Figure S1. Endogenous ZO-1 accumulated at the basal-front region of migrating cells**

Immunostaining images of the basal part of migrating MDCKII (parental).

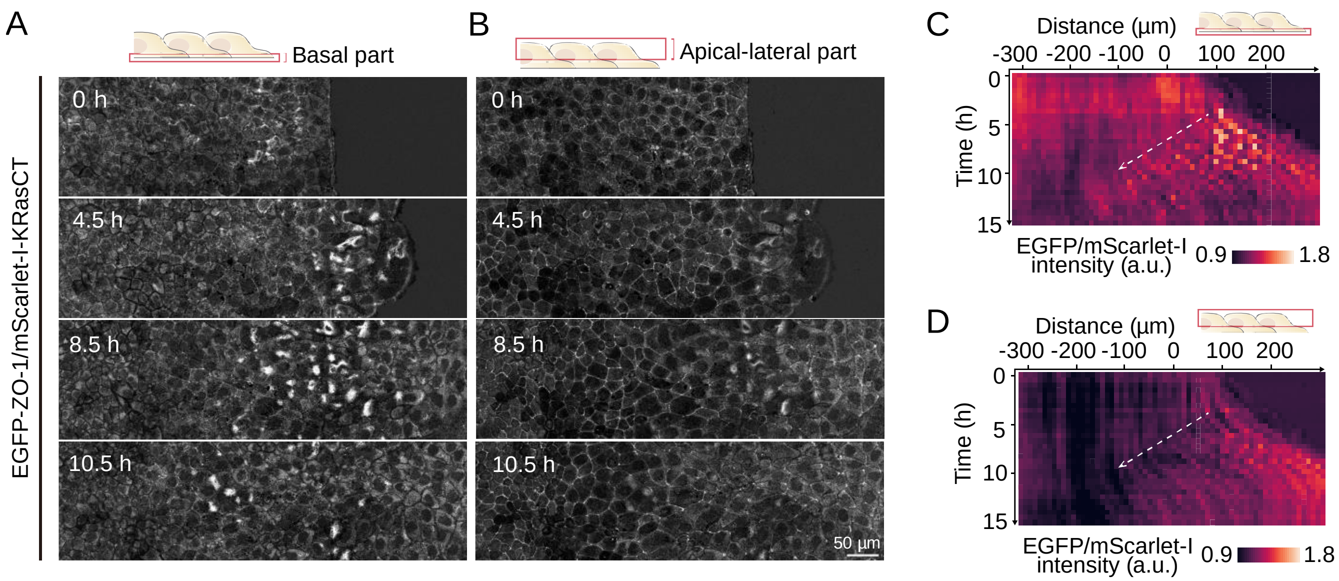


**Figure S2. ZO-1 translocation to the basal cell surface during collective cell migration**

(A, B) Timelapse images of the ratio of EGFP-ZO-1 to mScarlet-I-KRasCT. (A) and (B) show the basal part and the apical-lateral part of the same cell population, respectively.

(C, D) Kymographs of the average intensity of (A) and (B).

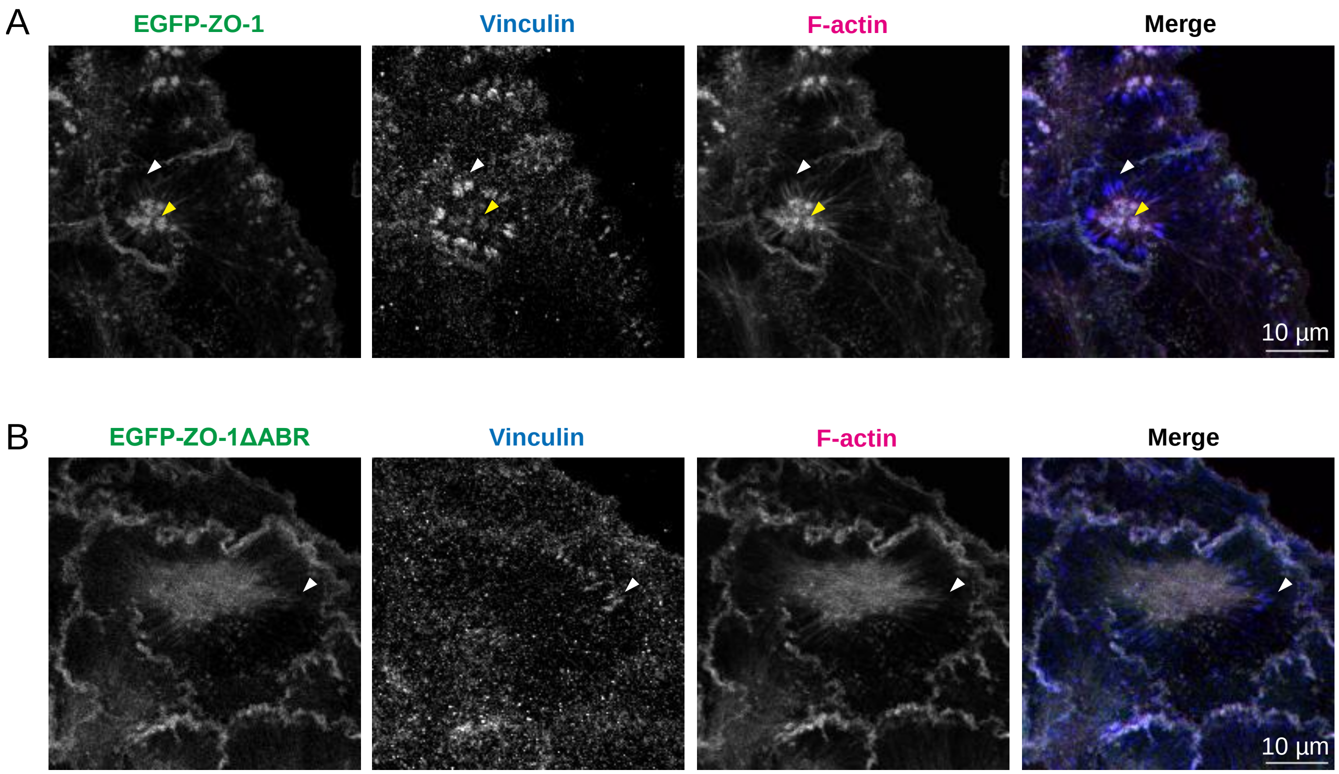


**Figure S3. Vinculin depletion from podosomes in ZO-1ΔABR-expressing cells**

(A, B) Immunostaining images of the MDCKII ZO-1/2dKO/EGFP-ZO-1 (A) and MDCKII ZO-1/2dKO/EGFP-ZO-1ΔABR (B) cell lines. Yellow and white arrowheads indicate podosomes and focal adhesions, respectively.

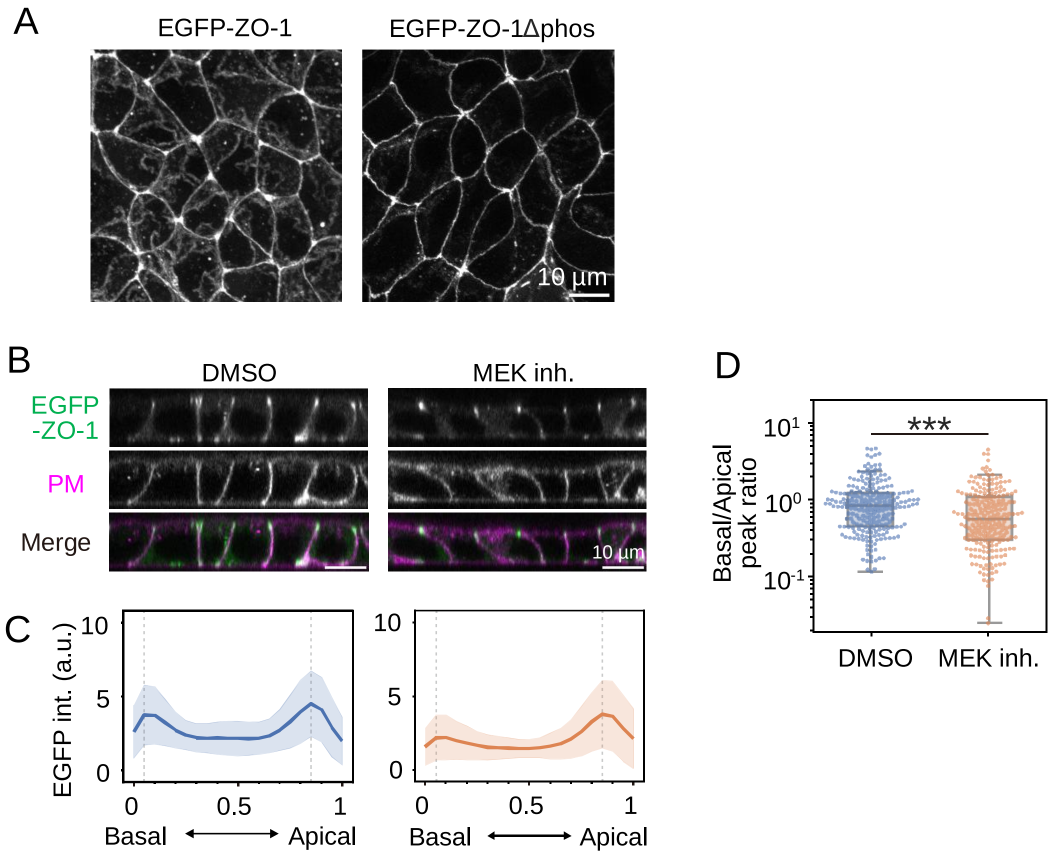


**Figure S4. Decrease in basal ZO-1 induced by inhibition of phosphorylation by ERK**

(A) Apical views of MDCKII ZO1/2dKO /EGFP-ZO1 and MDCKII ZO1/2dKO /EGFP-ZO1Δphos cells.

(B) Lateral views of the MDCKII ZO1/2dKO /EGFP-ZO1 cells treated with DMSO or 100 µM PD0325901 (MEKi) overnight. PM, plasma membrane.

(C) Quantified EGFP intensity along the lateral membranes. Data are from three independent experiments. The numbers of cell–cell interfaces were n = 227 (DMSO) and n = 214 (PD03). Lines and bands indicate the average and s.d., respectively.

(D) The ratio of EGFP intensity at the basal to apical peak positions. The peak positions are indicated in (C). Statistical significance was tested by the Mann–Whitney U-test.

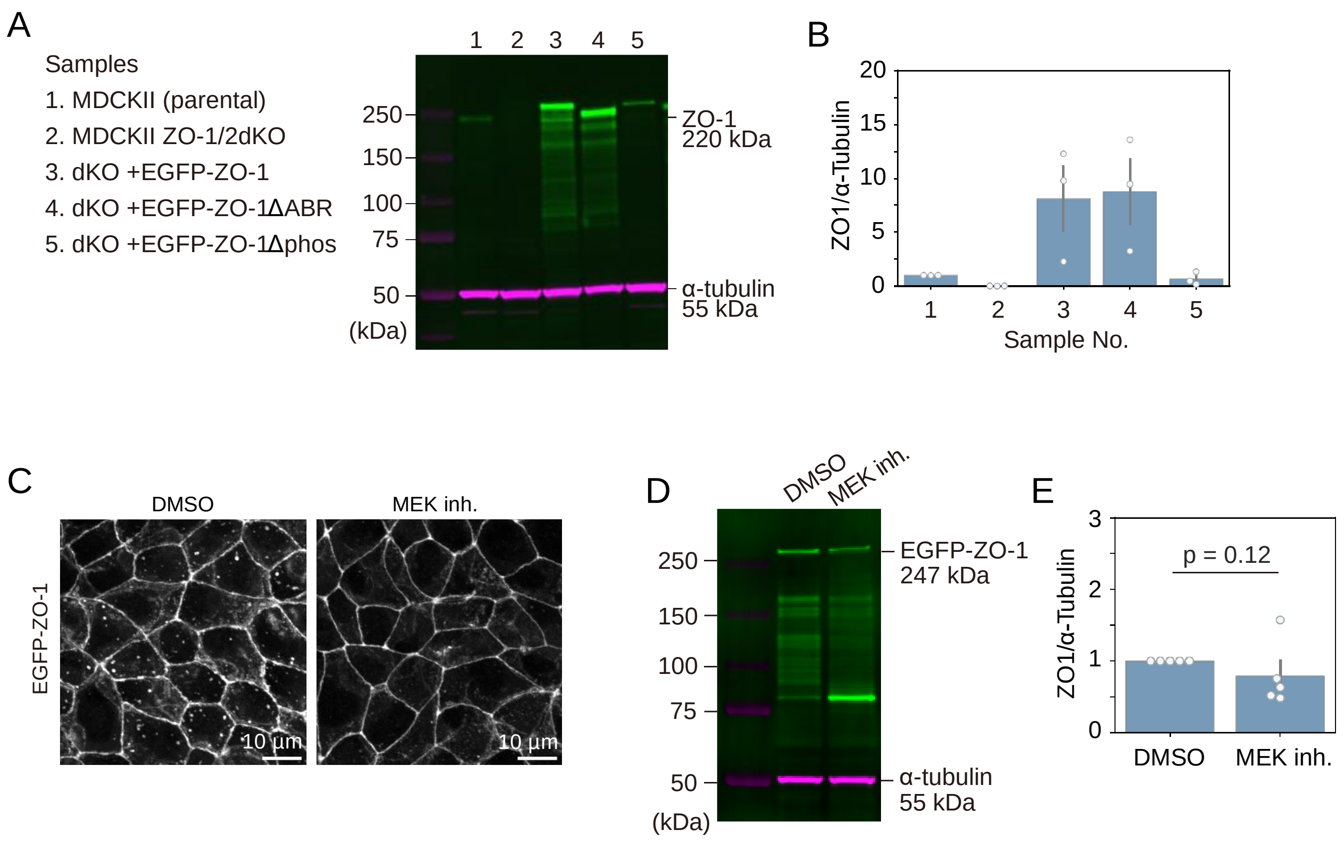


**Figure S5. Decrease in the level of ZO-1 proteins induced by inhibition of phosphorylation by ERK**

(A) Western blotting of ZO1 and alpha-tubulin for the stable cell lines.

(B) Quantified data of panel A. ZO1 was normalized with alpha-tubulin. The expression level in the parental MDCKII was set to 1. Data are from three independent experiments. Error bars indicate s.e.

(C) Apical views of the MDCKII ZO1/2dKO /EGFP-ZO1 cell line treated with DMSO or 100 μM PD0325901 (MEKi) overnight.

(D) Western blotting of ZO1 and alpha-tubulin for the cells treated with DMSO or 100 μM PD0325901 (MEKi) overnight.

(E) Quantified data of panel D. ZO1 was normalized with alpha-tubulin. The expression level in the cells treated with DMSO was set to 1. Data are from five independent experiments. Error bars indicate s.e. Statistical significance was tested by the Mann-Whitney U-test.

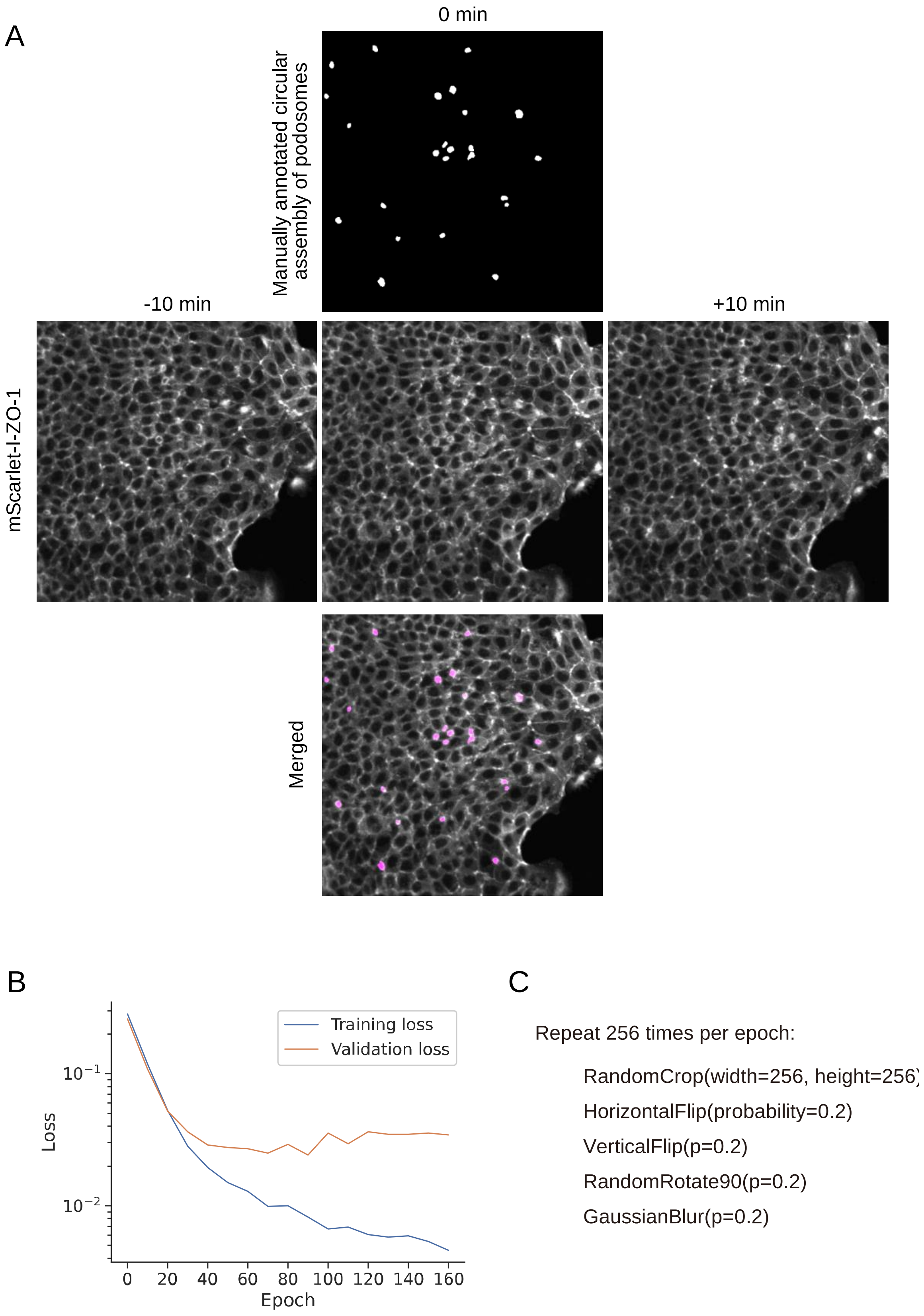


**Figure S6. Model training for automatic segmentation of the circular assemblies of podosomes**

(A) Representative training images for U-Net to detect circular assemblies of podosomes.

(B) The values of training loss and validation loss during the training.
(C) Pseudocode for data augmentation of the training dataset.

**Video S1. Translocalization wave of ZO-1 during collective cell migration, Related to Figure 1**
Collective migration of MDCK II cells that stably express EGFP-ZO-1 in the backbone of ZO-1/2 double KO. The apical part (upper panel) and the basal part (lower panel) of the same cell population are represented.

**Video S2. Circular assembly of podosome ZO-1 on the basal surface of migrating cells, Related to Figure 1**
Basal part of migrating MDCK II cells that stably express mScarlet-I-ZO-1 in the backbone of ZO-1/2 double KO cells.

**Video S3. ZO-1 accumulation to podosomes through actin-binding region, Related to Figure 2**
Basal part of MDCK II cells that stably expresses EGFP-ZO-1 (left and middle panels) or EGFP-ZO-1ΔABR (right panel) in the backbone of ZO-1/2 double KO cells. The cells were treated with DMSO or 10 nM TPA at the time point 0 minutes.

**Video S4. Formation of the circular assemblies of podosome ZO-1 following ERK activation during collective cell migration, Related to Figure 3**
Collective migration of MDCK II cells that stably express EKAREV-NLS (FRET-based ERK activity sensor) and mScarlet-I-ZO-1 in the backbone of ZO-1/2 double KO cells. The FRET/CFP ratio of EKAREV-NLS is represented in the intensity-modulated display mode with pseudo-colors. The time-lapse image of mScarlet-I-ZO-1 shows the basal surface of the cells.

**Video S5. Disappearance of podosome ZO-1 in response to MEKi, Related to Figure 4**
The basal part of MDCK II cells that stably express EGFP-ZO-1 in the backbone of ZO-1/2 double KO cells. The cells were treated with 10 nM TPA at the time point -60 min and further treated with DMSO or 100 µM PD0325901 (MEK inhibitor, MEKi) at the time point 0 minutes.

**Video S6. Reduced accumulation of ZO-1 in podosomes by non-phosphorylatable mutations, Related to Figure 4**

The basal part of MDCK II cells that stably express EGFP-ZO-1 (left panel) or EGFP-ZO-1Δphos (right panel) in the backbone of ZO-1/2 double KO cells. The cells were treated with DMSO or 10 nM TPA at the time point 0 minutes.

**Video S7. Intercellular propagation of ERK activation affected by the presence or absence of ZO-1 and mutations of ZO-1, Related to Figure 5**

ERK activity during collective cell migration visualized by EKEREV-NLS. The FRET/CFP ratio of EKAREV-NLS is represented in the intensity-modulated display mode with pseudo-colors. The cell lines are, from left to right, the parental MDCK II, MDCK II ZO-1/2 dKO, MDCK II ZO-1/2 dKO/EGFP-ZO-1, MDCK II ZO-1/2 dKO/EGFP-ZO-1ΔABR and MDCK II ZO-1/2 dKO/EGFP-ZO-1Δphos.
