## Supplementary material for "ZO-1 shuttles between tight junctions and podosomes by riding ERK activation waves during collective cell migration": Key resources table

| REAGENT or RESOURCE | SOURCE | IDENTIFIER |
|---|---|---|
| Antibodies | | |
| Mouse monoclonal anti-ZO-1 (ZO1-1A12) | Invitrogen | Cat#33-9100, Lot: YK381714 |
| Rabbit polyclonal anti-TKS5(SH3 #4) | Sigma-Aldrich | Cat#09-268, Lot: 4012809 |
| Rabbit polyclonal anti-Vinculin | Proteintech | Cat#26520-1-AP, Lot: 00079470 |
| Rabbit polyclonal anti-αTubulin | MBL | Ca#PM054, |
| Alexa Fluor Plus 488 conjugated goat polyclonal anti-mouse IgG (H+L) | Invitrogen | Ca# A32723 |
| Alexa Fluor Plus 405 conjugated goat polyclonal anti-rabbit IgG (H+L) | Invitrogen | Ca# A48254 |
| Alexa Fluor Plus 555 conjugated goat polyclonal anti-rabbit IgG (H+L) | Invitrogen | Ca# A32732 |
| IRDye 800CW Donkey anti-mouse IgG Secondary antibody | LI-COR | Ca#925-32212 |
| IRDye 680RD Goat anti-Rabbit IgG Secondary antibody | LI-COR | Ca# 926-68071 |
| Bacterial and virus strains | | |
| pCSIIpuro-EGFP-ZO1 | This study | https://benchling.com/s/seq-TpHCfJLP6s1B1O2yiEo0?m=slm-StLReL5ppAdsrYiPtLfD |
| pCSIIpuro-EGFP-ZO1ΔABR | This study | https://benchling.com/s/seq-fVHoGnpHDYEBPMWzfGQG?m=slm-aEAcHNBscCauL2OhaxYB |
| pCSIIpuro-EGFP-ZO1Δphos | This study | https://benchling.com/s/seq-7woM8xTBECAlGIntzacA?m=slm-EziGPp9v5FzRbwwzPajh |
| pCSIIbleo-H2B-iRFP | https://www.sciencedirect.com/science/article/pii/S1534580717308298?via=ihub - bib29 | https://benchling.com/s/cRQVQqr9?m=slm-g04dT7wmJCR2FNug6gzb |
| pCSIIhyg-mScarlet-I-KRasCT | This study | https://benchling.com/s/seq-xfCnEj5B0MAKj4G4zuH4?m=slm-x9g55nc5ZXFQBsUG8thn |
| Chemicals, peptides, and recombinant proteins | | |
| Alexa Fluor 633 Phalloidin | Invitrogen | Ca#A22284 |
| Phalloidin-iFluor 555 Conjugate | AAT Bioquest | Ca#23119 |
| CellMask Orange Plasma Membrane Stains | Invitrogen | Ca# C10045 |

| | | |
|---|---|---|
| TPA (PMA) | Sigma-Aldrich | Ca# P1585-1MG |
| PD0325901 | TOCRIS | Ca#4192 |
| Latrunculin A | Wako | Ca# 129-04361 |
| Experimental models: Cell lines | | |
| MDCKII | Gift from Mikio Furuse | N/A |
| MDCKII ZO1/2 double KO | Gift from Mikio Furuse | N/A |
| MDCKII ZO1/2 dKO/H2B-iRFP/EGFP-ZO1 | This study | N/A |
| MDCKII ZO1/2 dKO/H2B-iRFP/EGFP-ZO1ΔABR | This study | N/A |
| MDCKII ZO1/2 dKO/H2B-iRFP/EGFP-ZO1Δphos | This study | N/A |
| MDCKII/EKAREV-NLS | This study | N/A |
| MDCKII ZO1/2 dKO/EKAREV-NLS | This study | N/A |
| MDCKII ZO1/2 dKO/EKAREV-NLS/mScarlet-I-ZO1 | This study | N/A |
| MDCKII ZO1/2 dKO/EKAREV-NLS/mScarlet-I-ZO1ΔABR | This study | N/A |
| MDCKII ZO1/2 dKO/EKAREV-NLS/mScarlet-I-ZO1Δphos | This study | N/A |
| Human embryonic kidney cell 293-T (Lenti-X 293T cells) | Clontech | 632180 |
| Oligonucleotides | | |
| ZO-1_S167A_F GAGCTTGgCCCCGCGGTCAGACAGGCGG | This study | N/A |
| ZO-1_S167A_R CGCGGGGcCAAGCTCCTCTCTCTACTTGCAC | This study | N/A |
| ZO-1_S453A_F GAAGATgcCCCTGCAGCCAAGGAAGGCTTAG | This study | N/A |
| ZO-1_S453A_R TGCAGGGgcATCTTCTAGAACGCCAGCTACAAATATTC | This study | N/A |
| ZO-1_S530A_F AAAGGAAgCTCCCTATGGACTTAGTTTTAACAAAGG | This study | N/A |
| ZO-1_S530A_R TAGGGAGcTTCCTTTTCATATTCAAAATGGGTTCTAATATAGAAAG | This study | N/A |
| ZO-1_T708A_F AGATGTAgCACCAAATGCAGTTGATCGTCTTAAC | This study | N/A |
| ZO-1_T708A_R TTTGGTGcTACATCTAATAAAGCATGTTTGTCTTGATCTATG | This study | N/A |
| ZO-1_S1616A_F CCTGTGGcTCCTTCAGCTGTGGAAGAGG | This study | N/A |
| ZO-1_S1616A_R TGAAGGAgcCACAGGAATAGCTTTAGGCACTG | This study | N/A |
| ZO-1_S1694A_F CTGCTGgcTCCTTTGGTGATGTGTGGTCC | This study | N/A |
| ZO-1_S1694A_R CAAAGGAgcCAGCAGTGTTTCACCTTTCTCTTTATC | This study | N/A |
| Recombinant DNA | | |

| | | |
|---|---|---|
| pCS2-EGFP-ZO1 | https://www.sciencedirect.com/science/article/pii/S258900422200116X?via=ihub - sec4 | https://benchling.com/s/seq-V48Ni184y9h7vlUzylzF?m=slm-O7VedEATmti0wdx8vWEi |
| pCS2-EGFP-ZO1ΔABR | https://www.sciencedirect.com/science/article/pii/S258900422200116X?via=ihub - sec4 | https://benchling.com/s/seq-PeEBQmlo6KckEKJuYKFu?m=slm-pBNIuJfmKAn94CEY2XjG |
| pCSIIpuro-EGFP-ZO1 | This study | https://benchling.com/s/seq-TpHCfJLP6s1B1O2yiEo0?m=slm-StLReL5ppAdsrYiPtLfD |
| pCSIIpuro-EGFP-ZO1ΔABR | This study | https://benchling.com/s/seq-fVHoGnpHDYEBPMWzfGQG?m=slm-aEAcHNBscCauL2OhaxYB |
| pCSIIpuro-EGFP-ZO1Δphos | This study | https://benchling.com/s/seq-7woM8xTBECAlGIntzacA?m=slm-EziGPp9v5FzRbwwzPajh |
| pCSIIbleo-H2B-iRFP | https://www.sciencedirect.com/science/article/pii/S153458071730829 8?via=ihub - bib29 | https://benchling.com/s/cRQVQqr9?m=slm-g04dT7wmJCR2FNug6gzb |
| pCSIIhyg-mScarlet-I-KRasCT | This study | https://benchling.com/s/seq-xfCnEj5B0MAKj4G4zuH4?m=slm-x9g55nc5ZXFQBsUG8thn |
| psPax2 | gift from Didier Trono | Addgene Plasmid #12260 |
| pCMV-VSV-G-RSV-Rev | https://journals.asm.org/doi/full/10.1128/jvi.72.10.8150-8157.1998 | https://benchling.com/s/seq-dinqnagEmM43lGvo7gc7?m=slm-it7FigUveLba7MjjYjUD |
| pPBbsr2-EKAREV-NLS | https://www.molbiolcell.org/doi/full/10.1091/mbc.e11-01-0072 | https://benchling.com/s/seq-Qg04RTWZFV4ZotOaHVam?m=slm-ekwt8w3ONF0xrf2nig45 |

| | | |
|---|---|---|
| pT2Apuro-mScarlet-I-ZO1 | This study | https://benchling.com/s/seq-6CwrRPsgTXihAbLFiClW?m=slm-cafxeDKDyzTO5OulFIgS |
| pT2Apuro-mScarlet-I-ZO1ΔABR | This study | https://benchling.com/s/seq-KeIkHNqzg7yjEdTAPA0G?m=slm-OZtgSuTyfJAeEUbvl92p |
| pT2Apuro-mScarlet-I-ZO1Δphos | This study | https://benchling.com/s/seq-o95ktE4N2qZ2BCr5WpNJ?m=slm-RqEm3cayd3ibeZdXi9SF |
| pCAGGS-hyPBase | https://www.nature.com/articles/s41467-021-27458-3 - Sec8 | https://benchling.com/s/seq-oGkw53b41IZqvzF5yQ9K?m=slm-CfvvCEUOAGUr3jQm8b7w |
| pCAGGS-T2TP | gift of Koichi Kawakami | https://benchling.com/s/seq-yGo10mX966NnMTfpmt3F?m=slm-qMC2AU6Mr6ygjzN6hioZ |
| Software and algorithms | | |
| Fiji | https://www.nature.com/articles/nmeth.2019 | https://fiji.sc/ |
| Python version 3.9.12 | Python Software Foundation | https://www.python.org/ |
| LIM Tracker | https://www.nature.com/articles/s41598-022-06269-6 | https://github.com/LIMT34/LIM-Tracker |
| StarDist | https://link.springer.com/chapter/10.1007/978-3-030-00934-2_30 | https://github.com/stardist/stardist |
| Dense Optical Flow in OpenCV | https://link.springer.com/chapter/10.1007/3-540-45103-X_50 | https://github.com/opencv/opencv/blob/3.4/samples/python/tutorial_code/video/optical_flow/optical_flow_dense.py |
| Template Matching, PIV, FTTC | https://sites.google.com/site/qingzongtseng/imagejplugins?authuser=0 | https://sites.google.com/site/qingzongtseng/imagejplugins?authuser=0 |
| Pytorch-UNet | https://github.com/milesial/Pytorch-UNet | https://github.com/milesial/Pytorch-UNet |
